# PrimeKG-Plus: a refreshed and rare-disease-enriched precision medicine knowledge graph

**DOI:** 10.64898/2026.07.14.738415

**Authors:** Trinh Trung Duong Nguyen, Thuy Nguyen-Phuong, Quy-Hoai Nguyen, Amna Mumtaz Abbasi, Hanh-Dung Le Phan, Luong Bao-Anh Nguyen, Nhat-Thien Phan, Nurettin Nusret Curabaz, Alexander S. Hauser, Ziaurrehman Tanoli, Dinh Truong Nguyen, Albert J. Kooistra

## Abstract

**Background:** Disease-centered knowledge graphs (KGs) support drug repurposing and precision medicine research, yet many remain static after release while primary databases and literature continue to expand. PrimeKG (Precision Medicine Knowledge Graph) is a widely used multimodal KG providing a holistic view of diseases. However, its public release reflects a June 2021 data cutoff, omitting several years of subsequent data growth. This lag is especially consequential for rare diseases, where mechanistic and therapeutic evidence often remains scattered across publications rather than structured resources.

**Finding:** We present PrimeKG-Plus, a refreshed, rare-disease–enriched release of PrimeKG, rebuilt from all twenty original data resources updated to their December 2025 releases and three additional resources: OpenTargets, RepurposeDrugs, and nSIDES. Beyond synchronizing biomedical databases, PrimeKG-Plus captures approximately five years of previously unavailable rare-disease knowledge from the biomedical literature, curated from 637 PubMed abstracts and PubMed Central full-text articles using a language-model-assisted workflow focused on four rare neurological disorders: Canavan disease, Niemann–Pick disease type C, Tay–Sachs disease, and Batten disease. Extracted relations were refined through entity normalization, UMLS synonym mapping, embedding-based similarity ranking, and human expert review. Network topology analysis showed improved indirect drug–disease connectivity across three to six hops and added 447,288 drug–protein–disease paths linking previously unreachable drug–disease pairs. Temporal validation using drug approval records identified 55 molecular entities approved after the original PrimeKG June 2021 cutoff, 46 of which were absent from the original graph.

**Conclusion:** PrimeKG-Plus restores the temporal relevance of PrimeKG, providing an updated resource for drug repurposing, rare-disease research, and downstream machine-learning applications.

## Introduction

Developing a novel therapeutic typically requires 10–15 years and investments ranging from hundreds of millions to over US$1 billion, with high attrition rates throughout the development pipeline^1,2^. These conventional bottlenecks become overwhelmingly prohibitive when applied to rare diseases, which collectively affect approximately 300 million individuals globally with over 6000 distinct disorders^3^, might fail to offer commercial viability to offset such immense R&D risks. These challenges have motivated increasing interest in computational drug repurposing to identify new therapeutic indications for existing drugs with established safety profiles.

Recent advances in machine learning have further accelerated drug discovery and repurposing. In particular, biomedical knowledge graphs^4^ have emerged as a powerful framework for integrating heterogeneous biomedical data and enabling network-based computational analyses across diverse biomedical applications. Several large-scale resources have been developed to support this approach, including Hetionet^5^, DRKG^6^, CKC^7^, PanKB^8^, Bioteque^9^, PrimeKG^10^, RTX-KG2^11^, and ROBOKOP^12^. These integrative graphs connect diverse biomedical and clinical entities, enabling computational analyses that exploit network topology for applications such as drug–target prediction, network-based drug repurposing, adverse drug reaction prediction, disease mechanism discovery, and biomarker prioritization.

Among these resources, PrimeKG stands out as a comprehensive biological knowledge graph that integrates heterogeneous biomedical data within a disease-centered framework. By harmonizing curated information across 20 biological resources, spanning drug and disease ontologies, gene and protein annotations, pathways, phenotypes, anatomy, and molecular interaction networks, PrimeKG connects 10 entity types—gene/protein, drug, disease, anatomy, biological process, molecular function, cellular component, pathway, exposure, and effect/phenotype within a unified multi-relational schema comprising more than four million undirected relationships. This multi-scale, multi-relational structure has supported network-based drug repurposing^13,14^ and graph machine learning applications. It also suggests that PrimeKG has become a foundational resource for computational therapeutic discovery.

Despite its broad adoption and multimodal design, the current public release of PrimeKG reflects biomedical knowledge only through June 2021. Since then, major biomedical resources have expanded substantially. DrugBank^15^ and DrugCentral^16^ have released six and three major updates, respectively, while frequently updated resources such as the Comparative Toxicogenomics Database (CTD)^17^, Mondo Disease Ontology (MONDO)^18^, and Gene Ontology (GO)^19^ have each undergone more than 50 monthly releases. These heterogeneous update cycles illustrate the rapid evolution of biomedical knowledge.

However, synchronizing curated databases alone is insufficient. A substantial proportion of newly generated biomedical knowledge first appears in the primary literature and may take years to be incorporated into structured resources. This delay is particularly pronounced for rare diseases, where disease-specific molecular mechanisms, phenotypes, and therapeutic evidence are often reported in individual studies long before they become represented in reference databases. Consequently, literature curation remains essential for enriching biomedical knowledge graphs with emerging disease knowledge.

To address this limitation, we developed PrimeKG-Plus, an extended version of PrimeKG that expands this comprehensive framework along three complementary directions.

- First, we updated each of the original PrimeKG source databases to their latest available releases (through December 2025), allowing the integrated graph to reflect advances in curated biomedical databases since the original June 2021 cut-off.
- Second, we incorporated RepurposeDrugs^20^, OpenTargets^21^, and nSIDES^22^ databases to strengthen disease–drug associations, disease–gene relationships and therapeutic target evidence, and drug–adverse event associations.
- Third, we curated literature-derived relationships for four rare monogenic neurometabolic disorders^3^ of the central nervous system: Canavan disease (a metabolic leukodystrophy), and three lysosomal storage disorders (Niemann–Pick disease type C, Tay–Sachs disease and Batten disease). Together these disorders form a coherent rare-disease case-study set with overlapping neurometabolic mechanisms. They also represent precisely the setting in which literature-based curation adds the most value, because disease-specific evidence for such disorders is typically scattered across primary publications and under-represented in structured databases. At the same time, each has a substantial yet manageable body of published evidence, enabling rigorous expert manual curation across genes, disease mechanisms, phenotypes, and therapeutic associations. To ensure schema compatibility with the original knowledge graph, we used the relationship types predefined in PrimeKG.

These contributions position PrimeKG-Plus alongside, rather than in competition with, concurrent efforts to modernize biomedical knowledge integration, such as OptimusKG^23^, which assembles structured and semi-structured resources into a schema-constrained, ontology-grounded property graph. The two resources differ in emphasis: whereas OptimusKG re-architects the graph to maximize breadth and metadata richness across structured sources, PrimeKG-Plus preserves the original PrimeKG schema for direct compatibility with existing PrimeKG-based methods and benchmarks, and adds an expert-curated, literature-derived layer for rare diseases that structured-source integration alone does not capture.

These extensions were integrated into a single coherent resource, with all entities normalized to the Unified Medical Language System (UMLS)^24^ to provide standardized entity representation and facilitate interoperability with other biomedical databases and knowledge graphs.

## Methods

### Constructing the knowledge graph from up-to-date data releases

#### Overview of primary data resources

The following section describes the primary data resources used to construct PrimeKG-Plus (Figure 1). Data were retrieved using versioned snapshots where available; otherwise, the download dates are reported.

**Figure 1.**
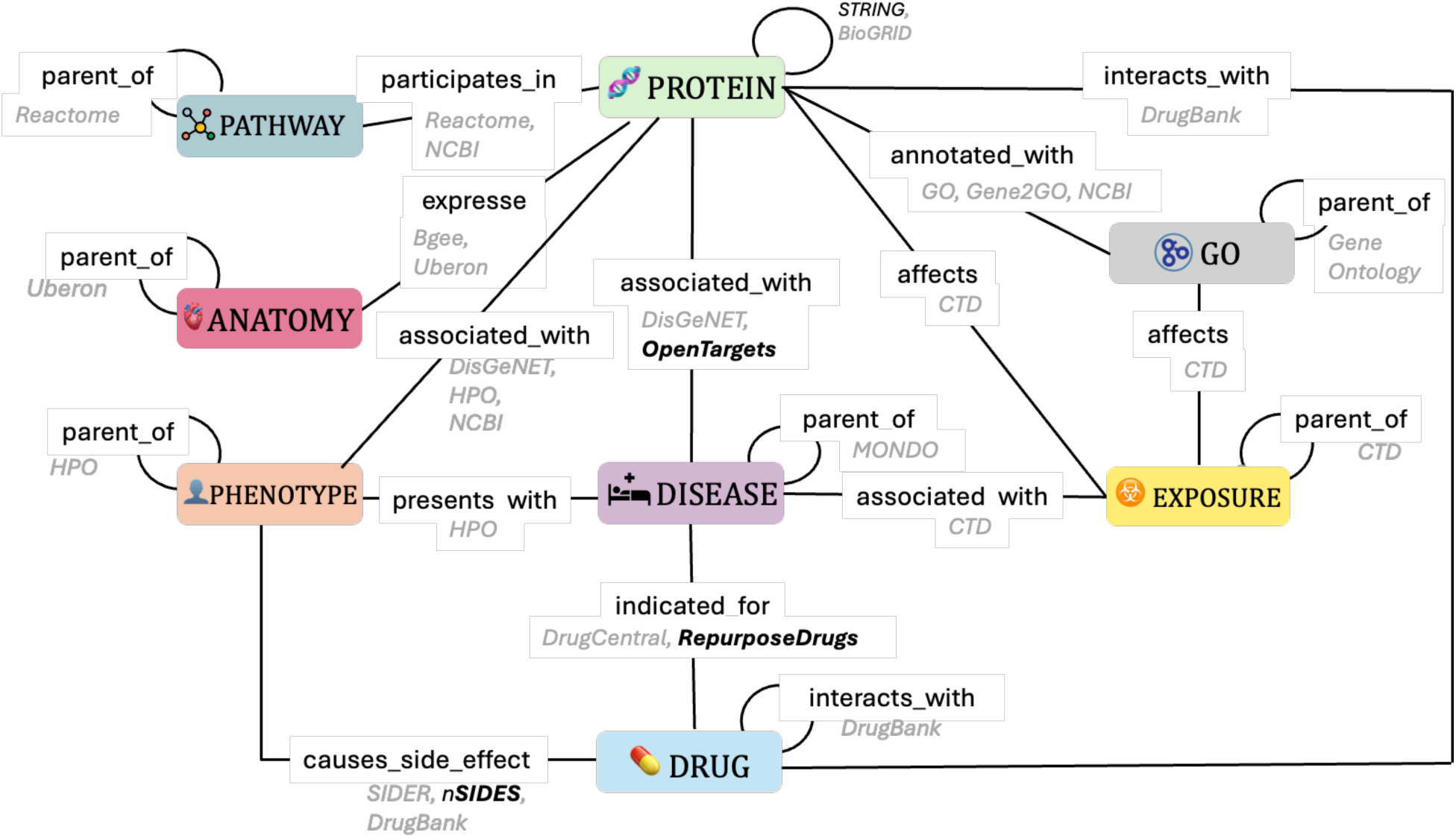
Construction of PrimeKG-Plus. PrimeKG-Plus extends PrimeKG by incorporating additional relationships from external resources. GO includes biological process (BP), cellular component (CC), and molecular function (MF). Edges denote relation types between entity categories, with data sources indicated alongside each edge; boldface database names indicate resources newly incorporated into PrimeKG-Plus.

Each relation type was rebuilt from updated primary-source snapshot using PrimeKG-compatible processing pipelines. Entity identifiers were harmonized across multiple ontologies, including MONDO, DrugBank, the Uber-anatomy ontology (UBERON)^25^, the Human Phenotype Ontology (HPO)^26^, GO, and others. Additional relation-specific quality-control procedures were implemented to improve data consistency and biological accuracy. These included MONDO disease-term filtering, bijective UMLS–MONDO mapping, HPO–MONDO disambiguation, and an updated Bgee expression-rank threshold for anatomy–protein associations. Following this, individual edge sets were concatenated, deduplicated, and augmented with reverse relations before extracting the largest connected component. Consequently, edge counts may increase or decrease relative to the original PrimeKG due to updates in source databases and the application of enhanced quality-control filters. Per-relation integration details are provided below.

Original PrimeKG served as the reference baseline for comparing edge counts, after removing 3,026 symmetric duplicate directed edges (0.04% of the directed edge set). Per-relation directed edge counts before and after integration are reported in Supplementary Tables S1–S2.

Edge-count convention. PrimeKG stores many biologically symmetric associations as directed edge records in both orientations (e.g., A→B and B→A for protein_protein, drug_drug, indication, contraindication, and drug_effect). Unless stated otherwise, all edge counts reported in this study are directed edge counts. Where helpful for interpretation, we additionally report unique unordered entity pairs (e.g., drug–drug and disease–gene pairs). Background references to PrimeKG’s “more than four million relationships” follow the original PrimeKG convention of counting undirected associations after collapsing reciprocal directed edges; our comparative analyses use directed counts for consistency with the released comma-separated values (CSV) edge lists and Supplementary Table S2.

### Elaboration on constructing different relation types

#### Protein-protein interactions

Protein-protein interactions were retrieved from the Search Tool for the Retrieval of Interacting Genes/Proteins (STRING) database (version 12.0) for human proteins (taxonomy ID 9606) and filtered to include only highest-confidence interactions^27^ with a combined confidence score ≥ 900. The STRING protein identifiers were mapped to Entrez Gene IDs^28^ by merging with a Swiss-Prot^29^ dataset containing reviewed human proteins, where entries with missing STRING identifiers or Gene IDs were excluded, and Gene IDs with multiple values were filtered out to ensure one-to-one mapping. The resulting interactions were represented as pairwise Gene ID relationships and deduplicated. Cross-referencing against the original PrimeKG (642,150 directed protein–protein edges across 18,354 proteins) identified 123,570 novel directed edges involving 338 additional proteins. After integration and deduplication, PrimeKG-Plus contains 765,720 directed protein– protein edges (+19.2% vs original PrimeKG).

### Drug-protein interactions

DrugBank is a curated resource containing pharmaceutical knowledge. We retrieved DrugBank version 5.1.13 from the official release portal on 29 November 2025 and processed the dataset using three scripts provided by the PrimeKG authors to maintain compatibility with the original schema. The updated release increases directed drug–protein interactions across all interaction categories in the final graph, including carriers (proteins carrying drugs; 2,108 vs 1,728), enzymes (drug-metabolizing proteins; 12,400 vs 10,634), targets (proteins responsible for therapeutic drug effects; 41,360 vs 32,544), and transporters (membrane proteins mediating drug transport; 7,080 vs 6,030), yielding 62,948 directed drug–protein edges in PrimeKG-Plus (vs 50,936 in original PrimeKG; +23.6%).

#### Protein-disease associations

In the original PrimeKG study, protein–disease associations were derived from DisGeNET^30^ (Disease–Gene Network). Because the legacy DisGeNET download interface was unavailable at the time of our analysis, we obtained the curated dataset directly from the PrimeKG authors and retained only disease-type associations, excluding group-level and phenotype-type entries (the latter are described under Phenotype–protein associations). To increase coverage, we incorporated additional gene–disease associations from OpenTargets (version 25.12.0), retaining direct target– disease associations with overall score > 0.1 (justification provided in Supplementary Information, Section C). Associations from both sources were harmonized to MONDO ontology using the curated bijective UMLS–MONDO vocabulary (see Supplementary Information, Section D) and merged with disease–protein edges retained from original PrimeKG. The updated DisGeNET processing table contained 84,038 curated gene–disease association rows, and OpenTargets contributed 75,361 additional disease–protein association records after MONDO harmonization. In the final graph, disease–protein relations increased from 160,813 to 516,740 directed edges (+221.3%), corresponding to 80,411 vs 258,262 unique disease–gene pairs (Supplementary Table S2).

### Drug-disease relations

Following the approach used by the PrimeKG authors, we retrieved the DrugCentral database dump (released on 10 May 2023) and extracted drug–disease relationships, including indications, contraindications, and off-label uses. We further incorporated approved indication edges from RepurposeDrugs (Phase 4), adding 383 novel indication pairs after removing 65 pairs already present in DrugCentral. After MONDO mapping and deduplication at the (drug, disease, relation) level, PrimeKG-Plus contains 10,006 indication edges, 26,826 contraindication edges, and 2,398 off-label use edges (vs 18,776, 61,350, and 5,136 in original PrimeKG). Decreases relative to original PrimeKG primarily reflect stricter MONDO disease-term filtering, UMLS-to-MONDO mapping with single Concept Unique Identifier (CUI) resolution (n_umls = 1), and removal of ambiguous drug–disease links during the updated DrugCentral integration. All harmonization used the curated vocabulary, described below and in Supplementary Information, Section D, rather than the raw cross-reference table.

### Filtering disease terms from MONDO

All datasets containing disease-related information (e.g., protein–disease associations, drug– disease relations, and exposure–disease associations) were harmonized using the MONDO disease ontology to standardize disease identifiers. We retrieved the MONDO release dated 2025-11-04 and used it as the primary disease vocabulary. Although MONDO integrates disease concepts from multiple biomedical ontologies, the raw ontology file also contains non-disease terms such as chemical entities, biological processes, anatomical structures, and gene symbols, arising from extensive cross-referencing to ChEMBL^31^ (Chemical database of EMBL), GO, UBERON, and the HUGO Gene Nomenclature Committee (HGNC)^32^. No explicit filtering step for such terms was described in the original PrimeKG documentation and scripts. To maintain the disease-centric structure of PrimeKG-Plus, we therefore retained only terms explicitly classified as diseases, as indicated by disease-specific subsets (e.g., mondo_disease, ordo_group, mondo_rare) or by explicit disease definitions within MONDO, excluding all terms lacking disease classification or exclusively cross-referenced to non-disease ontologies.

### Drug–drug interactions

Synergistic drug–drug interactions were extracted from DrugBank (version 5.1.13, downloaded 29 November 2025) using the DrugBank vocabulary (17,430 drugs, downloaded 24 November 2025). We parsed all drug-interaction entries in the DrugBank Extensible Markup Language (XML) release to obtain directed DrugBank-ID pairs, retained only pairs in which both drugs mapped to the vocabulary, and exported them as drug–drug edges, following the original PrimeKG schema. This yielded 2,855,310 directed drug–drug edges among 4,566 drugs (1,427,655 unique unordered pairs), compared with 2,672,628 directed edges in original PrimeKG (+182,682; +6.8%). drug_drug is the most frequent relation type in PrimeKG-Plus (37.2% of directed edges; Supplementary Table S2).

### UMLS–MONDO Vocabulary Curation

PrimeKG-Plus uses a bijective UMLS–MONDO vocabulary, where UMLS concepts are represented by Concept Unique Identifiers (CUIs), to prevent spurious edge duplication during graph assembly. The original PrimeKG vocabulary was constructed to maximize cross-source coverage by concatenating all MONDO–UMLS cross-references without resolving one-to-many associations, causing ambiguous CUI-to-MONDO or MONDO-to-CUI mappings to propagate as duplicate edges through inner joins. PrimeKG-Plus instead applies explicit bijective curation, trading broader mapping recall for edge-level precision.

The raw vocabulary contained 31,683 pairs across 22,418 unique MONDO IDs, with 20.2% of MONDO IDs mapping to more than one UMLS CUI and 10.4% of UMLS CUIs claimed by multiple MONDO concepts. We queried the Monarch Initiative application programming interface (API) for unambiguous UMLS cross-references; for remaining ambiguous cases, we queried the UMLS Terminology Services API and selected the CUI with the highest preference score, prioritizing preferred terms (PT; termType = PT) from SNOMED CT US Edition (SNOMEDCT_US), Medical Subject Headings (MSH), National Cancer Institute Thesaurus (NCI), Online Mendelian Inheritance in Man (OMIM), and Orphanet Rare Disease Ontology (ORDO) (ties discarded). The final curated vocabulary contained 21,357 bijective MONDO– UMLS pairs and was used for all disease harmonization steps in PrimeKG-Plus. Full multiplicity analysis, pipeline replay, and worked examples are provided in Supplementary Information, Section D (Supplementary Tables S10–S11).

### Anatomy-protein associations

Human gene expression calls were retrieved from the Bgee database^33^ (release 2025-02-03). Anatomical entities were mapped to UBERON identifiers, retaining only gold quality expression calls annotated with presence/absence and expression rank. The updated Bgee release has finer anatomical granularity, which inflated edge counts under the original PrimeKG cutoff (expression rank < 25,000; lower ranks indicate higher expression). We therefore applied a stricter threshold (rank ≤ 5,000) to preserve high-expression signals while keeping a comparable association density. After filtering, the dataset contributed 930,465 unique anatomy-protein present pairs (1,860,930 directed edges), compared with 3,033,804 directed edges in original PrimeKG (-38.7%; Supplementary Table S2).

### Adverse-drug effects

Adverse-drug effect data were processed in two steps. First, we reproduced the author Side Effect Resource (SIDER) pipeline (SIDER 4.132): records mapped from MedDRA (Medical Dictionary for Regulatory Activities; https://www.meddra.org/) restricted to Preferred Terms were joined with drug–Anatomical Therapeutic Chemical (ATC) mappings on compound identifiers and output as one row per adverse-drug effect pair; the MedDRA-linked UMLS CUI was used as the canonical adverse-effect identifier and duplicate (drug, adverse effect) pairs were removed. Second, by adding nSIDES data, we augmented this table with high-confidence adverse-drug effect associations from Open nSIDES. RxNorm ingredient identifiers were mapped to ATC codes using DrugBank, and nSIDES MedDRA effect names were mapped to UMLS CUIs by matching the SIDER side-effect vocabulary. Only pairs with valid ATC mapping were retained. Finally, in PrimeKG-Plus, drug_effect edges total 121,840 directed relations (vs 129,568 in original PrimeKG; −6.0%).

### Exposure related associations

Comparative Toxicogenomics Database (CTD) exposure events (downloaded October 2025) were processed with header cleanup and MONDO mapping for disease-related edges. Exposure– protein, exposure–disease, exposure–GO, and exposure–exposure relations were rebuilt from the updated CTD release. The final graph contains 4,770 exposure–disease edges (+3.5%), 6,086 exposure–protein edges (+151.1%), and 4,206 exposure–biological process edges (+29.4%) compared with original PrimeKG (Supplementary Table S2).

### Other relation types

The remaining hierarchical and annotation relations were rebuilt from updated source releases using the PrimeKG processing pipeline: disease–disease parent–child relations (MONDO, release 2025-11-04); disease–phenotype (positive and negative) and phenotype–phenotype hierarchies (HPO, release 2025-10-22); phenotype–protein associations from DisGeNET phenotype-type records with HPO/MONDO disambiguation; protein–GO annotations and GO term hierarchies (GO release 2025-10-10; updated National Center for Biotechnology Information (NCBI) gene2go table); pathway–protein and pathway–pathway relations (Reactome); and anatomy– anatomy hierarchies (UBERON snapshot 2025-11-24). Pairs in which one phenotype mapped to a MONDO disease concept were converted to disease_phenotype_positive relations, consistent with the disease-centric PrimeKG schema. Disease grouping and subtype consolidation are described in the disease-grouping section. Detailed per-source processing steps are provided in Supplementary Information - Part I.

Edge counts cited in the main text are derived from processed source files prior to final graph integration. The released knowledge graph may contain marginally fewer edges after node-identifier mapping, deduplication, self-loop removal, and extraction of the largest connected component.

### Literature-Derived Relationship Extraction

#### Literature retrieval

We retrieved abstracts from PubMed^34^, which served as the primary input for relation extraction using the E-utilities API^35,36^. For open-access records linked to PubMed Central (PMC), full-text articles were also retrieved and used selectively during expert validation, with curators consulting Introduction or Conclusion sections when abstract-level evidence was insufficient to confirm or disambiguate a relation. For each disease, the search query followed a common structure: one MeSH (Medical Subject Headings) term for the disease plus optional Title/Abstract (TIAB) terms for name variants and for the main enzyme or gene deficiency. All terms were combined with OR and passed as the ‘term’ parameter; results were restricted by publication date (mindate, maxdate). The approach was kept strict (MeSH + selected TIAB terms only) to limit false positives.

Applied to the four rare neurological disorders:

- **Canavan disease**: MeSH "Canavan Disease"; TIAB variants "Canavan disease", "Canavan’s disease"; deficiency terms "ACY2 deficiency", "Aspa deficiency", "Aspartoacylase deficiency".
- **Tay–Sachs disease**: MeSH "Tay-Sachs Disease"; TIAB variants "Tay-Sachs disease", "Tay Sachs disease", "Sachs disease"; deficiency/classification terms "Hexosaminidase A deficiency", "HEXA deficiency", "GM2 Gangliosidosis Type 1", "GM2 gangliosidosis type 1".
- **Niemann–Pick disease type C**: MeSH "Niemann-Pick Disease, Type C"; TIAB variants "Niemann-Pick disease type C", "Niemann-Pick type C", "NPC disease"; deficiency terms "NPC1 deficiency", "NPC2 deficiency".
- **Batten disease** (neuronal ceroid lipofuscinosis): MeSH "Neuronal Ceroid Lipofuscinoses"; TIAB variants "Batten disease", "neuronal ceroid lipofuscinosis", "NCL"; optionally gene/type terms such as "CLN1", "CLN2", "CLN3" as needed.

Date filters (e.g., from 2021/06/01 or 2021/06/21 to 2025/11/06) were applied per disease; retrieval was capped (e.g., retmax = 10,000). **Figure 3** shows the number of retrieved articles per disease.

### Artificial intelligence (AI)-assisted relation extraction and expert validation

In this workflow, each journal article serves as the input text, and predefined curation requirements act as a structured prompt. The prompt specifies a fixed set of 30 relationship types aligned with PrimeKG (e.g., protein–protein, drug–protein, disease–protein, drug–disease indication, phenotype associations, pathway interactions, and anatomy-related relations) to ensure structural consistency with the reference knowledge graph. The same prompt template was applied uniformly across all articles to standardize entity and relation extraction; the complete prompt specification is provided in Supplementary Information, Section A, with the supported relation types enumerated in Supplementary Table S6.

Both the article and the prompt were integrated into NotebookLM to generate preliminary entity– entity relations along with associated metadata (entity types, relation types, and source identifiers). The extracted results were subsequently cross verified against the articles and manually curated by medical students to remove false positives, resolve taxonomic ambiguities, and preserve biological context. All validated relationships underwent an additional review by a PhD-level domain expert to ensure biological plausibility and semantic accuracy (see more on Supplementary Information, section B). NotebookLM (Google) was used between December 2025 and March 2026 with the latest Gemini model available within the platform at that time (Gemini 2.5 Flash or Gemini 3). Generation parameters (e.g., temperature) are not exposed to users in the closed-platform environment and were therefore not controlled. The prompt template and the list of curated articles are provided in the associated Github repository to support reproducibility. The complete curation workflow, including curator roles, validation procedures, and quality-control steps, is illustrated in Supplementary Figures S1 and S2.

### Entity harmonization of curated relations

We developed a three-stage workflow (**Figure 2**) to harmonize and reassess curated biomedical relations for integration into PrimeKG-Plus and applied this workflow to four understudied diseases. An entity was considered present if its normalized name matched an existing PrimeKG entity name. We first identified entity names that were not already represented in PrimeKG. These novel entities were then processed through a three-stage normalization workflow consisting of (i) UMLS-based normalization and validation, (ii) semantic recovery of unmapped entities using PrimeKG-guided retrieval, and (iii) a second UMLS lookup and relation filtering.

i. UMLS-based normalization and validation: Entity names were first normalized using rule-based methods (abbreviation expansion, typo correction, and canonical naming), then mapped to UMLS via the Representational State Transfer (REST) API (accessed March 2026). For each normalized term, the first valid concept match was selected based on lexical similarity, returning a Concept Unique Identifier (CUI) along with its semantic type. Every concept has a semantic type which is also associated with an identifier, the TUI (Type Unique Identifier). Matches were considered valid only when at least one retrieved TUI overlapped with the predefined expected type group for that entity category (detailed in Table S9, Section B, Supplementary Information); otherwise, they were flagged as semantically inconsistent. The resulting CUIs, matched names, and TUI annotations were propagated back to both endpoints of each relation pair, and only relations with valid mappings on both ends were marked as high-confidence records.
ii. Semantic recovery of unmapped entities using PrimeKG-guided retrieval: Entities that failed the initial UMLS validation were reassessed using embedding-based semantic matching. Each unmapped entity was compared against the PrimeKG entity vocabulary using SapBERT^37^ embeddings (Self-alignment Pretrained Bidirectional Encoder Representations from Transformers [BERT]), and the top 20 candidates ranked by cosine similarity were reranked using a Sentence-BERT (SBERT)^38^ model to select the best replacement. This dual-embedding design leverages complementary strengths: SapBERT provides strong biomedical concept alignment through UMLS-based supervision, while SBERT improves contextual discrimination among semantically similar candidates. For example, the term "enzyme enhancement" was reassigned to "enzyme regulator activity", yielding a valid UMLS mapping (CUI C1152521; Molecular Function). Selected replacements were accepted only after a second ontology-based validation step confirming semantic type compatibility, and validated records from both stages were merged into a unified output table for downstream integration.
iii. Second UMLS lookup and relation filtering: Each SBERT-selected replacement was re-queried against UMLS and retained only if the returned CUI matched the expected semantic type for that PrimeKG relation. Accepted replacements were propagated to their original entity strings, and each endpoint was assigned one of three statuses: “in_kg” (exact match to an existing PrimeKG node label), a UMLS CUI, or invalid. Relations were kept only when both endpoints received a non-invalid status. This automated pass yielded 1,087 integratable triples across four diseases (98 Canavan; 193 Batten; 729 NPC; 67 Tay-Sachs). Entity names that remained unmapped after this pass were exported as a manual-review pool (1,109 unique entity names; 25, 437, 596, and 51 by disease) for downstream quality control (QC) review.

**Figure 2.**
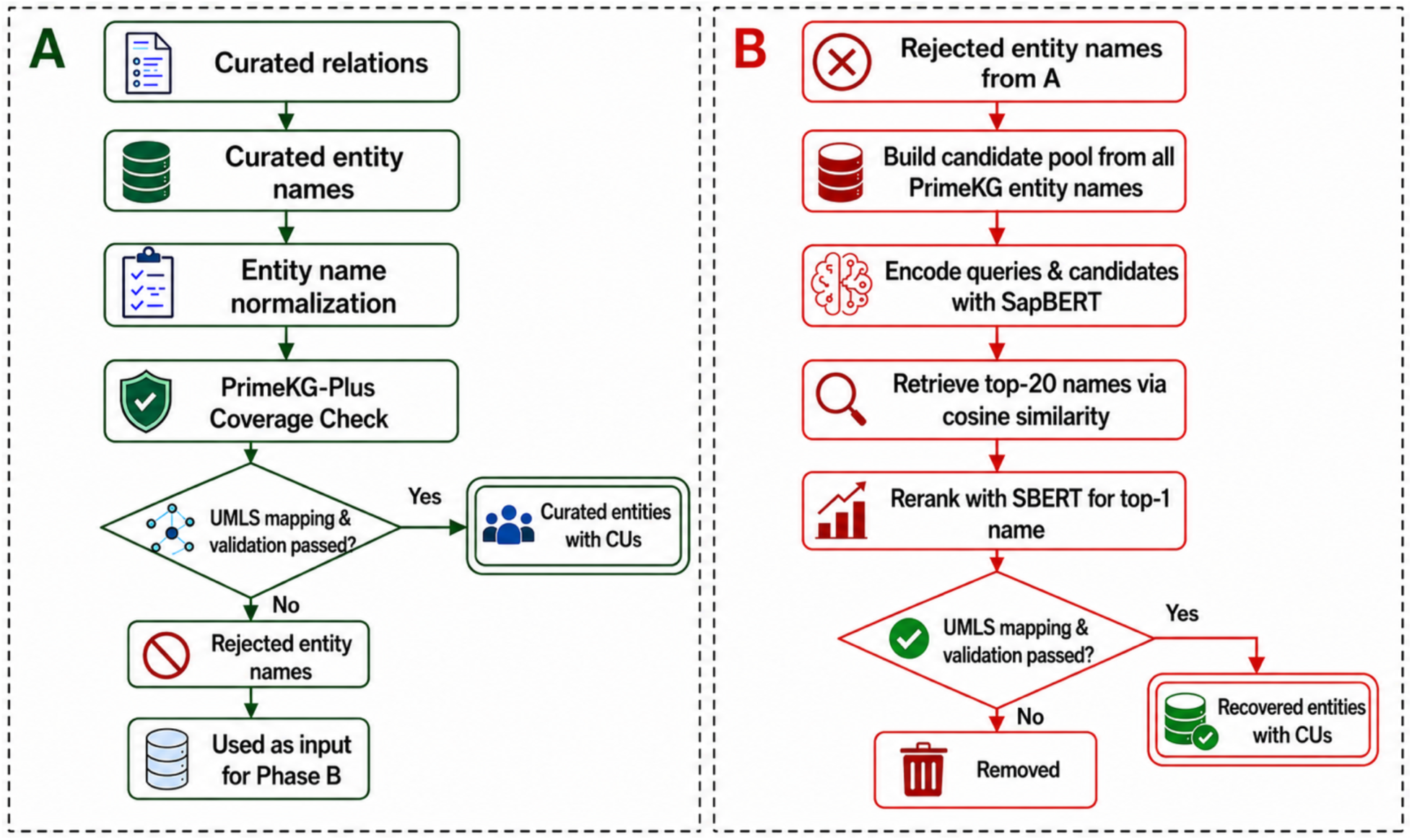
Biomedical entity curation and recovery workflow. **(A)** Curated entity names extracted from literature-derived relations are first normalized and compared against existing PrimeKG entities. Entity names already present in PrimeKG are retained directly, whereas only novel entity names are subjected to UMLS mapping and validation. Novel entities that are successfully mapped to and validated against UMLS concepts are retained as curated entities with CUIs, whereas entities that fail validation are retained as rejected entity names. **(B)** Rejected entity names from phase A are recovered through semantic retrieval over PrimeKG entity names using SapBERT embeddings, the top 20 candidates are retrieved based on cosine similarity, and the highest-ranked candidate is selected through SBERT-based re-ranking. Recovered entities are subsequently subjected to UMLS mapping and validation. Entities passing validation are retained as recovered entities with CUIs, whereas the remaining entities are discarded.

The final output of this workflow are UMLS-validated entities with CUIs (double line boxes, Figure 2) that can be linked to PrimeKG-Plus nodes and used for literature-edge integration.

### Second-round expert entity review

Entity names that the automated pipeline could not resolve were compiled into a manual-review pool. Subsequently, medical experts cross-referenced this list directly against the original article context to eliminate false positives, address missing entities, and correct any misclassifications, ensuring quality control. The detailed workflow is provided in the Supplementary Fig. S1-2. For each pooled entity, reviewers also saw whether the automated second-UMLS pass on the BERT-suggested replacement had already produced an acceptable mapping. QC team members and domain experts annotated preferred concept labels and UMLS CUIs where available. Expert-approved identifiers were merged back into the full curated corpus, re-evaluating all relation triples under the expanded mapping and recovering additional integratable relations beyond the algorithmic finals. Per-disease funnel counts are reported in Figure 3.together with the algorithmic final relation tables for context

**Figure 3.**
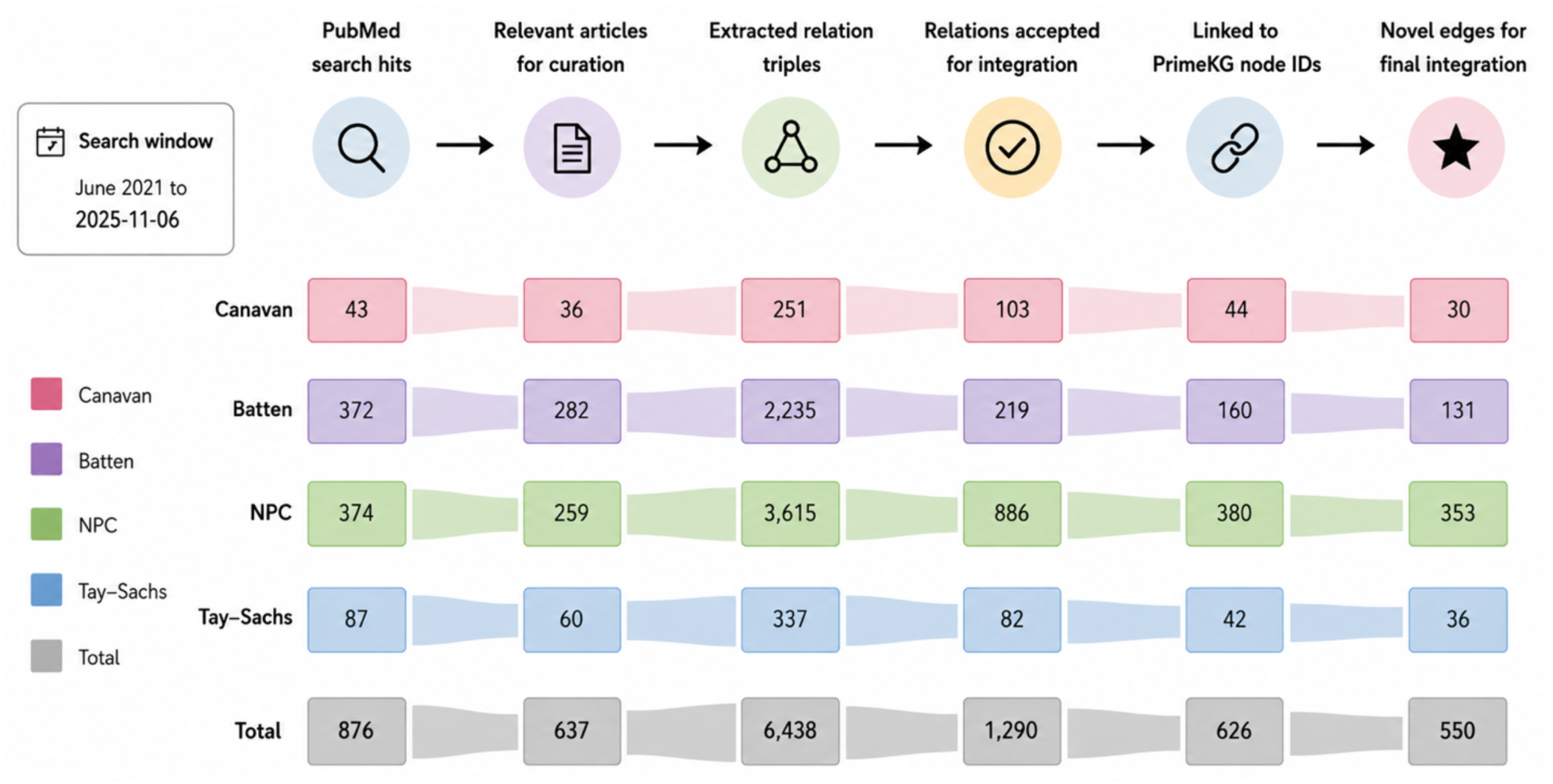
Literature curation funnel for four rare neurological disorders.

### Graph integration of curated literature relations

Each accepted relation was linked to PrimeKG-Plus by assigning both endpoints a graph node identifier from the released node inventory. Harmonized entity strings were matched to existing node labels where possible; otherwise, UMLS CUIs from the curation pipeline were mapped to ontology identifiers (MONDO, HPO, NCBI, DrugBank, GO, UBERON) using PrimeKG cross-reference tables and UMLS Metathesaurus atom names. When a validated concept corresponded to a canonical ontology term not yet represented in the graph, a new literature-specific node was created. Curated rows follow the PrimeKG-compatible relation schema summarized in Supplementary Table S8, with worked examples in Supplementary Table S7; supported relation types are listed in Supplementary Table S6. Relations that could not be integrated most often failed because at least one endpoint—despite passing harmonization and frequently carrying a UMLS CUI, could not be mapped to any resolvable ontology atom in PrimeKG-Plus. Other exclusions were duplicate literature rows or relation types outside the PrimeKG schema. Integration outcomes by disease are summarized in Figure 3.

### Data Description

PrimeKG-Plus is released as two PrimeKG-compatible knowledge-graph builds in CSV format, together with companion node and edge tables, literature-curation outputs, and auxiliary rebuild files. Both builds follow the original 12-column PrimeKG edge-list schema, enabling drop-in reuse with existing PrimeKG analysis workflows.

The base build, primekg_plus.csv, integrates updated public biomedical resources through December 2025—including Open Targets, RepurposeDrugs, and nSIDES—and contains no PubMed-derived edges. The literature-augmented build, primekg_plus_rd.csv, contains every edge from the base build plus expert-validated literature-derived relations for four rare neurological disorders: Canavan disease, Niemann–Pick disease type C, Tay–Sachs disease, and Batten disease. The public directory layout of the release bundle is shown in Figure 4. Raw licensed upstream dumps are not redistributed and can be retrieved from the original providers when a full rebuild is required.

**Figure 4.**
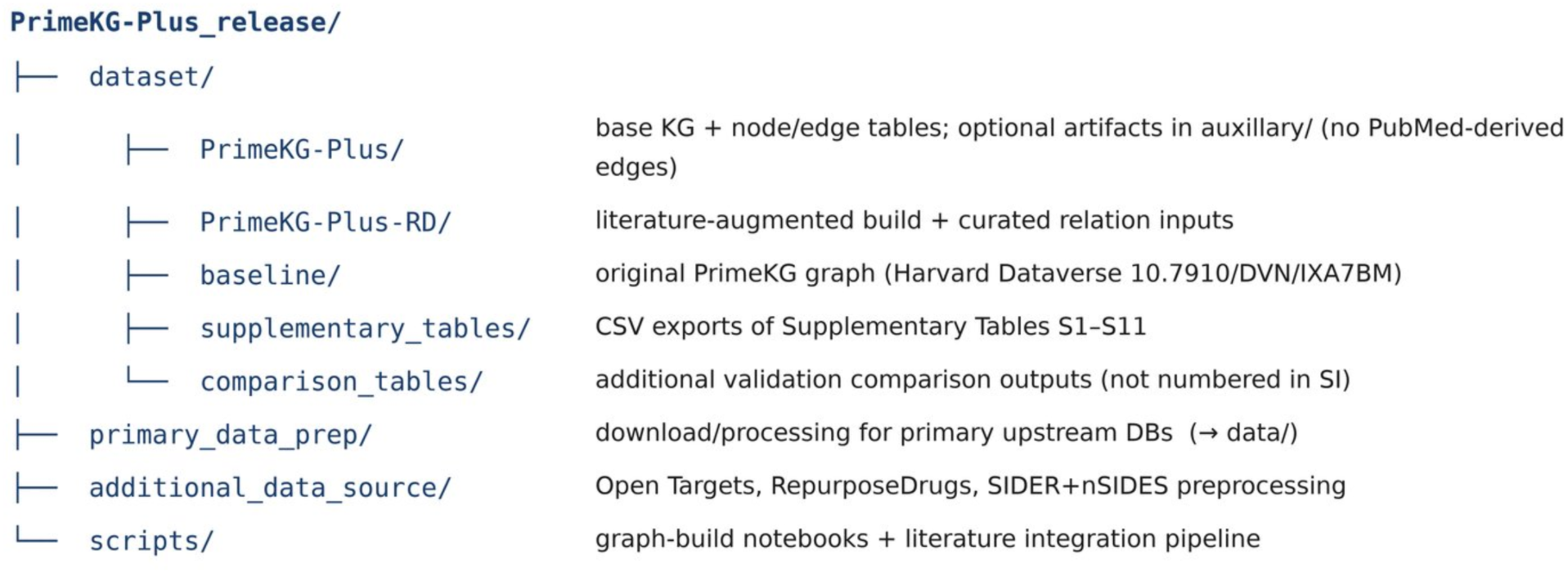
Directory layout of the PrimeKG-Plus release bundle.

The numbers of articles and extracted relations in the literature integration workflow are summarized in **Figure 3**: **(1) Literature retrieval.** Disease-specific PubMed searches from June 2021 onward returned 876 records across the four diseases (43 Canavan; 372 Batten; 374 NPC; 87 Tay–Sachs). **(2) Article selection for curation.** After screening, 637 unique articles contained extractable biomedical relations (36; 282; 259; 60 by disease). **(3) Relation extraction.** The curation team extracted 6,438 raw entity–relation triples from these articles (251; 2,235; 3,615; 337). **(4) Entity harmonization and validation.** Entity names were mapped to PrimeKG-compatible identifiers (PrimeKG labels, UMLS CUIs, and ontology IDs) and reviewed by domain experts; duplicate triples across diseases were removed. This yielded 1,290 relations accepted for integration (103; 219; 886; 82). **(5) Graph linking**. Each accepted relation was resolved to PrimeKG node identifiers; 626 relations were successfully linked (44; 160; 380; 42). **(6) Graph integration.** Of the 626 linked relations, 550 were novel relative to primekg_plus.csv and were added to primekg_plus_rd.csv (30; 131; 353; 36), together with 36 new ontology nodes required to represent previously unmapped literature entities. The remaining linked relations were already present in the base graph. Relations not integrated (664 rows in total) included duplicates, unsupported relation types outside the PrimeKG schema, or endpoints that could not be mapped to graph nodes.

Together, these products define the PrimeKG-Plus data records evaluated in the following validation analyses.

### Data Validation and quality control

This section presents technical validation and quality control for PrimeKG-Plus. We summarize graph-composition changes relative to Original PrimeKG, assess disease-node harmonization, and evaluate the literature-extraction workflow that underpins the curated rare-disease edges. Temporal updating, multi-hop drug–disease connectivity, and rare-disease neighborhood enrichment are illustrated as reuse-oriented case studies in the Re-use potential section.

#### Coverage Comparison

PrimeKG-Plus contains 129,317 entities versus 129,375 in Original PrimeKG (−0.04%). Disease-centered entities expanded (disease +16.4%; effect/phenotype +22.1%; drug +17.9%), whereas gene/protein (−14.5%) and biological process (−12.2%) decreased following updated harmonization and ontology pruning. Disease and effect/phenotype shares rose to 15.4% and 14.5%, respectively (Figure 5; Supplementary Table S1).

**Figure 5:**
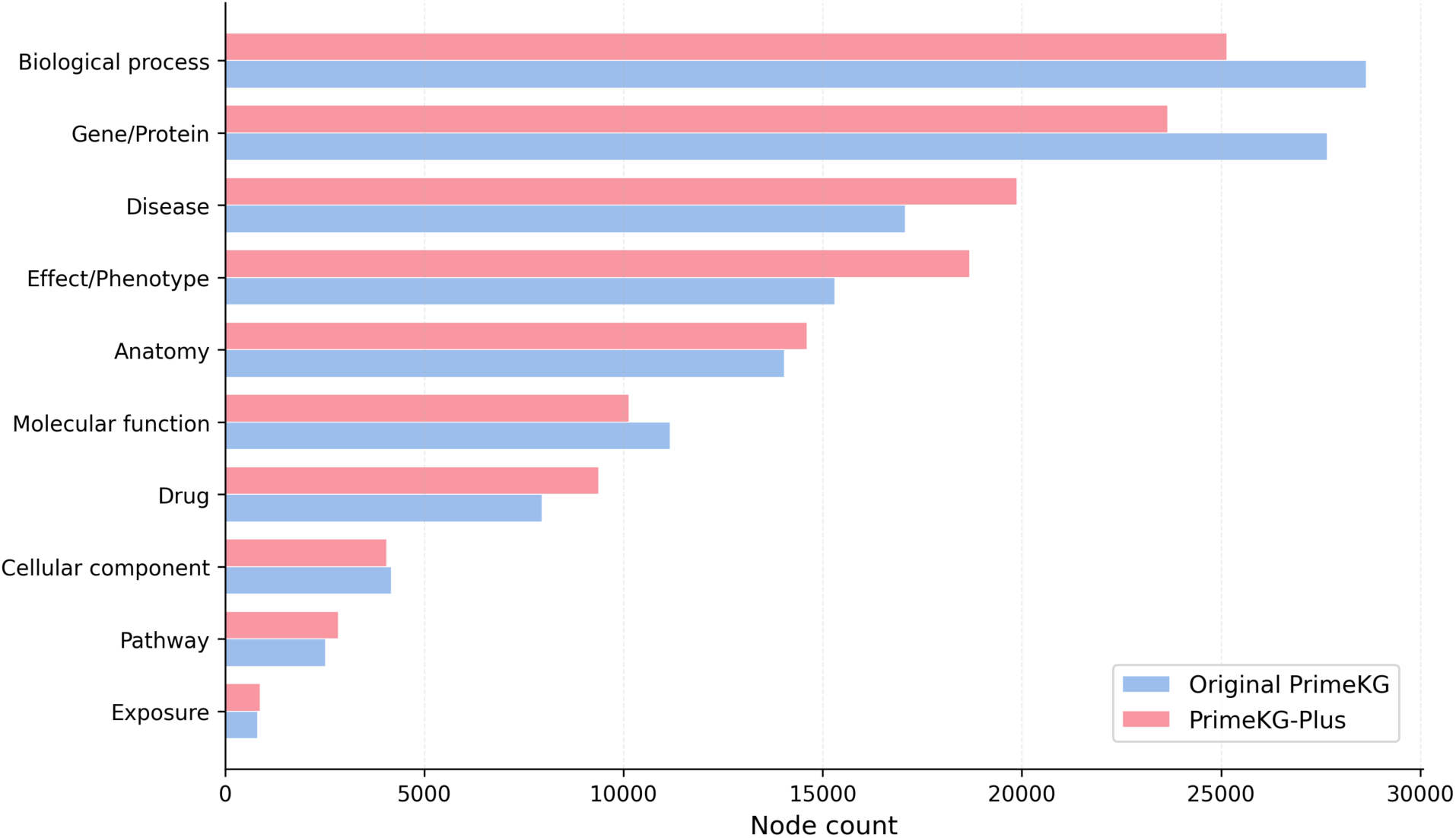
Entity type distribution in original PrimeKG and PrimeKG-Plus.

Directed edges decreased by 5.1% (8.10M → 7.68M). The largest drop was anatomy_protein_present (−38.7%; Bgee rank filter), whereas disease_protein increased 3.2-fold (+221%) and disease_phenotype_positive, protein_protein, and drug_drug also rose. Drug–disease edges declined under stricter MONDO curation, and drug_drug became the most frequent relation type (37.2% vs 24.2%) (Figure 6; Supplementary Table S2). Overall, PrimeKG-Plus does not uniformly increase edge counts; instead, it reallocates graph density toward disease–gene and phenotype knowledge while reducing redundant anatomy-expression and ambiguous drug–disease links.

**Figure 6:**
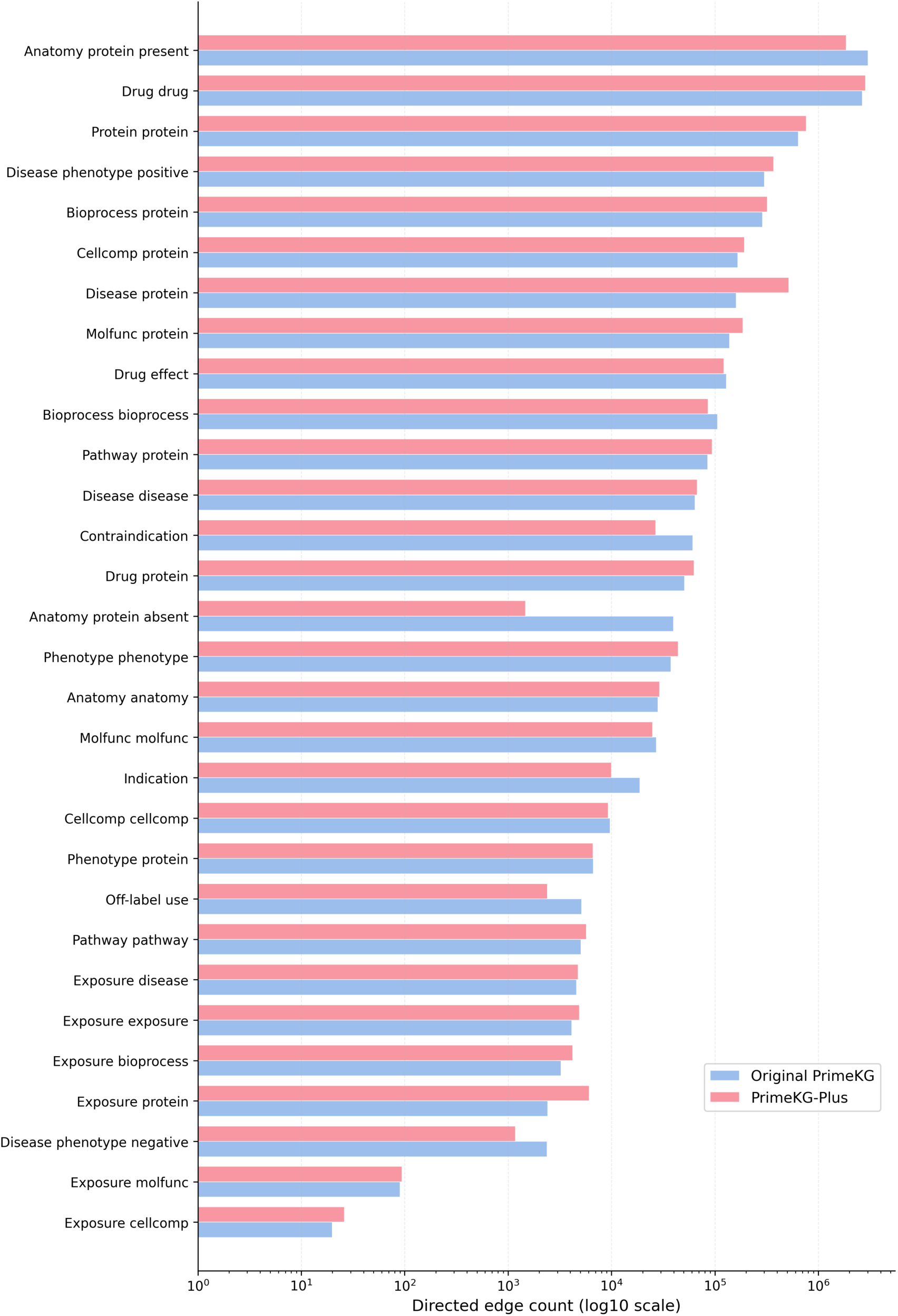
Distribution of relationship types (in terms of directed edges) in original PrimeKG and PrimeKG-Plus.

#### Disease node harmonization and grouping

Disease nodes were filtered to MONDO disease terms (release 2025-11-04) and grouped into MONDO_grouped hub nodes using the same workflow as original PrimeKG (rule-based checks, BERT-based semantic similarity, and lookup against published PrimeKG groupings). Groups reproducing an original PrimeKG hub were kept as-is; newly formed or modified groups underwent domain expert review before integration. This consolidation reduces sparsely connected disease singletons whose weak neighborhoods can degrade graph embedding quality and downstream link prediction.

PrimeKG-Plus contains 18,704 singleton MONDO disease nodes and 1,180 MONDO_grouped hubs, compared with 15,813 MONDO and 1,267 MONDO_grouped nodes in Original PrimeKG (Supplementary Table S3). Mean incident degree on grouped hubs rose to 214.4 (vs 115.1), reflecting edge aggregation onto hubs (Supplementary Table S4). Quality control identified 865 disease names shared between graphs that map to different node identities; comparisons should therefore use (node_id, node_type, node_source) rather than display names (node_name). Both graphs remain >99.9% contained within a single connected component (99.95% and 99.91% of nodes in Original PrimeKG and PrimeKG-Plus, respectively).

### Literature extraction evaluation

The literature extraction workflow used a structured prompt of approximately 800 tokens specifying 30 PrimeKG-compatible relationship types, well within the context capacity of the Gemini models (Gemini 2.5 Flash, later upgraded to Gemini 3 with approximately 1 million tokens) used within NotebookLM, allowing both prompt and article text to be processed without truncation. The complete prompt template is provided in Supplementary Information, Section A, and the supported relation types are listed in Supplementary Table S6.

When processing large numbers of articles within a single NotebookLM session, we occasionally observed a gradual decrease in extraction throughput. This behaviour is consistent with previously reported “positional bias” effects in large language models, where model performance degrades when relevant information is farther to the end of the input. To mitigate this effect, literature extraction was performed in smaller batches, and the extraction environment was periodically refreshed to maintain stable performance. Minor variability in entity naming across repeated extractions (e.g., abbreviation substitution, plural forms) was resolved during the normalization and expert validation stages.

To ensure high data quality, the curation process followed a structured human-in-the-loop workflow. Curators used a standardized extraction template that records the extracted relation together with the source location in the article (e.g., abstract, introduction, or conclusion); output fields are defined in Supplementary Information, Section A, Supplementary Table S5 (literature curation output columns), and Supplementary Table S8 (column definitions for curated relationships), with representative rows in Supplementary Table S7. Each curated relation was cross-checked by another curator and subsequently reviewed by a domain expert, enabling traceability of the evidence and ensuring consistency across the curated dataset.

### Re-use potential

#### Data reuse for studies in life sciences research

PrimeKG-Plus is designed to support a broad range of AI-driven biomedical research tasks, particularly those requiring a large-scale, multi-relational knowledge graph as a structured backbone. The original PrimeKG graph has been widely adopted as a benchmark resource with applications spanning drug–disease indication and contraindication prediction^14^, drug–target interaction^39^, drug–drug interaction prediction^40^, link prediction across heterogeneous biomedical entity types^39^, and zero-shot therapeutic inference for diseases lacking known treatments^13^. PrimeKG-Plus extends this foundation with updated database versions and literature-derived edges for four rare neurological disorders, enabling more accurate and temporally current benchmarking of these tasks. The updated disease–protein associations and expanded drug coverage further support protein–disease association mining and network-based drug repurposing pipelines.

Beyond graph machine learning benchmarks, the final knowledge graph files (primekg_plus.csv and primekg_plus_rd.csv) can be readily converted into related relational databases^41^, as each triple can be represented as a relational record linking two entities through a defined relationship, facilitating integration with Structured Query Language (SQL)-based infrastructure or graph databases. The dataset is also compatible with agentic AI platforms such as ToolUniverse^42^, which provides a standardized interface for connecting large language models to scientific databases and tools, enabling autonomous biomedical research workflows.

In addition, the dataset is designed in accordance with the FAIR (Findable, Accessible, Interoperable, and Reusable) principles. Researchers are free to use the dataset provided that the present study is appropriately cited.

### Case studies

The following case studies illustrate concrete reuse scenarios for the released graphs, complementing the technical validation above.

#### CS1: Temporal coverage of post-2021 FDA NMEs

Case study 1 asks which U.S. Food and Drug Administration (FDA) Type 1 – New Molecular Entity (NME) approvals after the Original PrimeKG snapshot (June 2021) are represented in PrimeKG-Plus. We compiled FDA approval records from July 2021 through December 2025 and retained Type 1 – NME submissions with Approval status (n = 149 approval rows; 143 unique active ingredients). Each active ingredient was matched to drug nodes in PrimeKG-Plus by exact case-insensitive string identity. Fifty-five (55) post-cutoff NME active ingredients were represented as drug nodes in PrimeKG-Plus; of these, 46 correspond to drug nodes absent from Original PrimeKG, whereas 9 (including Elafibranor, Ganaxolone, Maribavir, Omidenepag isopropyl, Sotagliflozin, Sparsentan, Tapinarof, Tirzepatide, and Treosulfan) were already present in the original release. The remaining 88 NME ingredients were not matched under exact-name rules, largely because FDA labels include salt or formulation suffixes whereas PrimeKG-Plus stores base drug names.

#### CS2: Drug–disease connectivity after graph update

Case study 2 examines whether the update changes multi-hop connectivity for drug–disease pairs that lack a direct indication edge. We compared the proportion of such pairs reachable within k-hop shortest paths (k = 2–6) in PrimeKG and the PrimeKG-Plus full graph (Figure 7). Analyses used 104,738,152 drug–disease pairs from the shared drug (7,845) and disease (13,352) node sets that lacked a direct indication edge in both Original PrimeKG and PrimeKG-Plus. As shown in **Figure 7**, PrimeKG-Plus showed higher reachability at k = 3–6. For example, the fraction of reachable pairs increased from 33.09% to 39.28% for k = 3, from 75.69% to 85.78% for k = 4. At k = 2, reachability was slightly lower in PrimeKG-Plus (4.26% vs 5.08%; Δ = −0.82). These results indicate that disease-centric edge enrichment, particularly disease–protein expansion, improves multi-hop connectivity for non-indicated drug–disease pairs without disrupting global graph connectivity. We treat this as a structural sanity check for multi-hop repurposing rather than a direct performance metric; downstream predictive benchmarking was outside the scope of this resource update.

**Figure 7.**
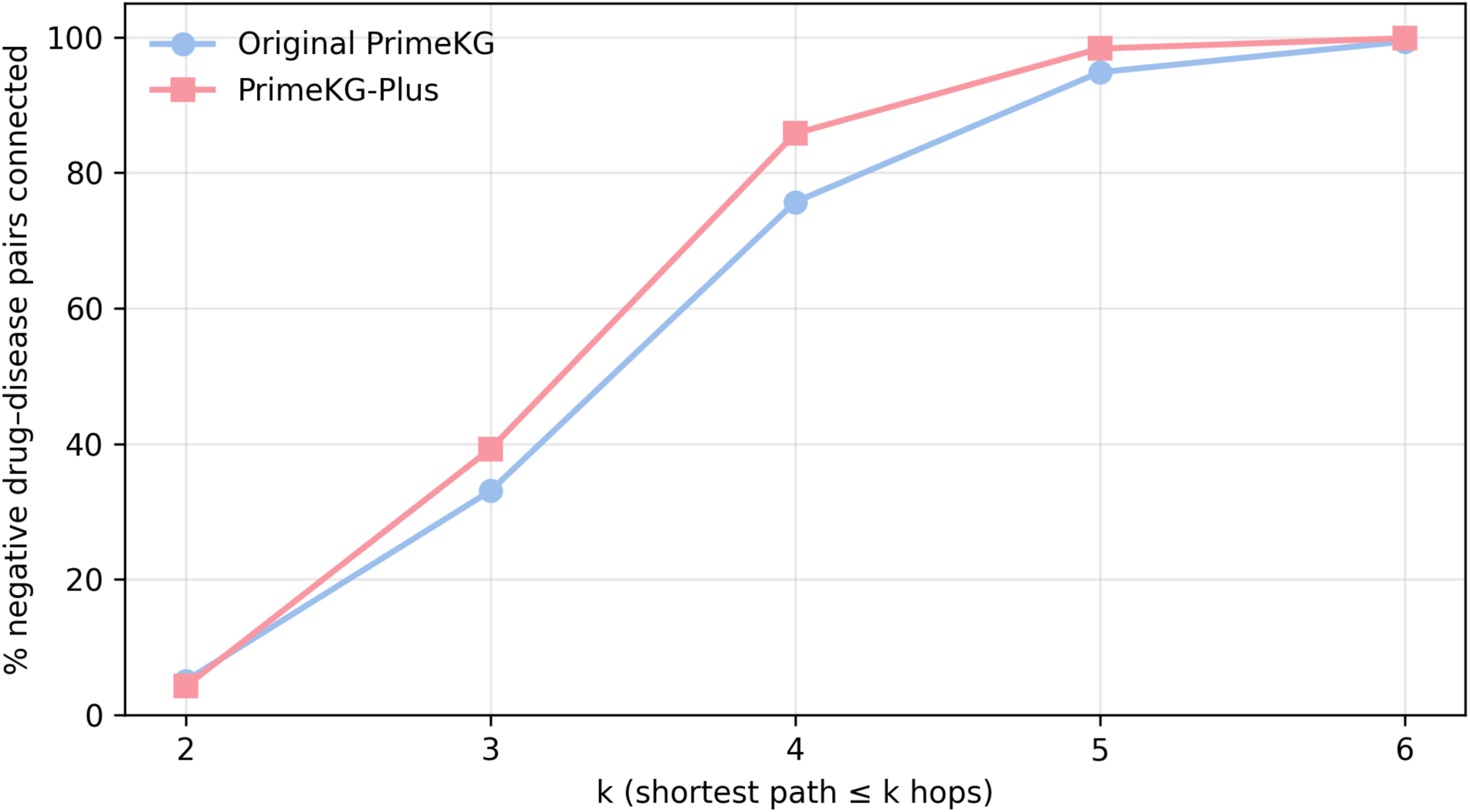
Drug–disease connectivity in negative pairs (no direct indication in either graph)

To illustrate mechanistic reuse beyond aggregate reachability, we counted Drug→Protein→Disease paths present in PrimeKG-Plus but absent from Original PrimeKG, keeping only those whose endpoint drug and disease could not be connected within six hops in the legacy graph. This yielded 447,288 interpretable two-hop drug–target–disease chains that the published graph could not recover. This count refers to specific protein-bridged paths for previously disconnected pairs, not to the fraction of all drug–disease pairs reachable within two hops (Figure 7). Two drugs approved after the Original PrimeKG snapshot illustrate this update. Avacopan (DrugBank DB15011) is a C5a receptor 1 (C5AR1) antagonist indicated for anti-neutrophil cytoplasmic antibody–associated vasculitis. Original PrimeKG retained the disease node (Microscopic polyangiitis) but lacked Avacopan entirely, so no route linked drug, target, and disease. PrimeKG-Plus adds Avacopan, connects it to C5AR1 and microscopic polyangiitis, and records the approved indication. Daridorexant (DrugBank DB15031), a dual orexin receptor antagonist approved in 2022 for insomnia, was also missing from the legacy release; PrimeKG-Plus links Daridorexant to insomnia via HCRTR1 and HCRTR2 together with a direct indication edge, which are absent from Original PrimeKG. These cases show that synchronizing drug, protein target, disease, and indication in this update restores mechanistic explanations for post-cut-off therapies that a static knowledge graph cannot provide.

#### CS3: Literature-enriched rare-disease neighborhoods

Case study 3 focuses on reuse around the four literature-curated rare neurological disorders. For each disease, we defined a disease scope comprising the primary anchor node plus named subtypes and synonyms and counted novel edges and short paths relative to Original PrimeKG. Summary statistics are reported in Figure 8.

**Figure 8.**
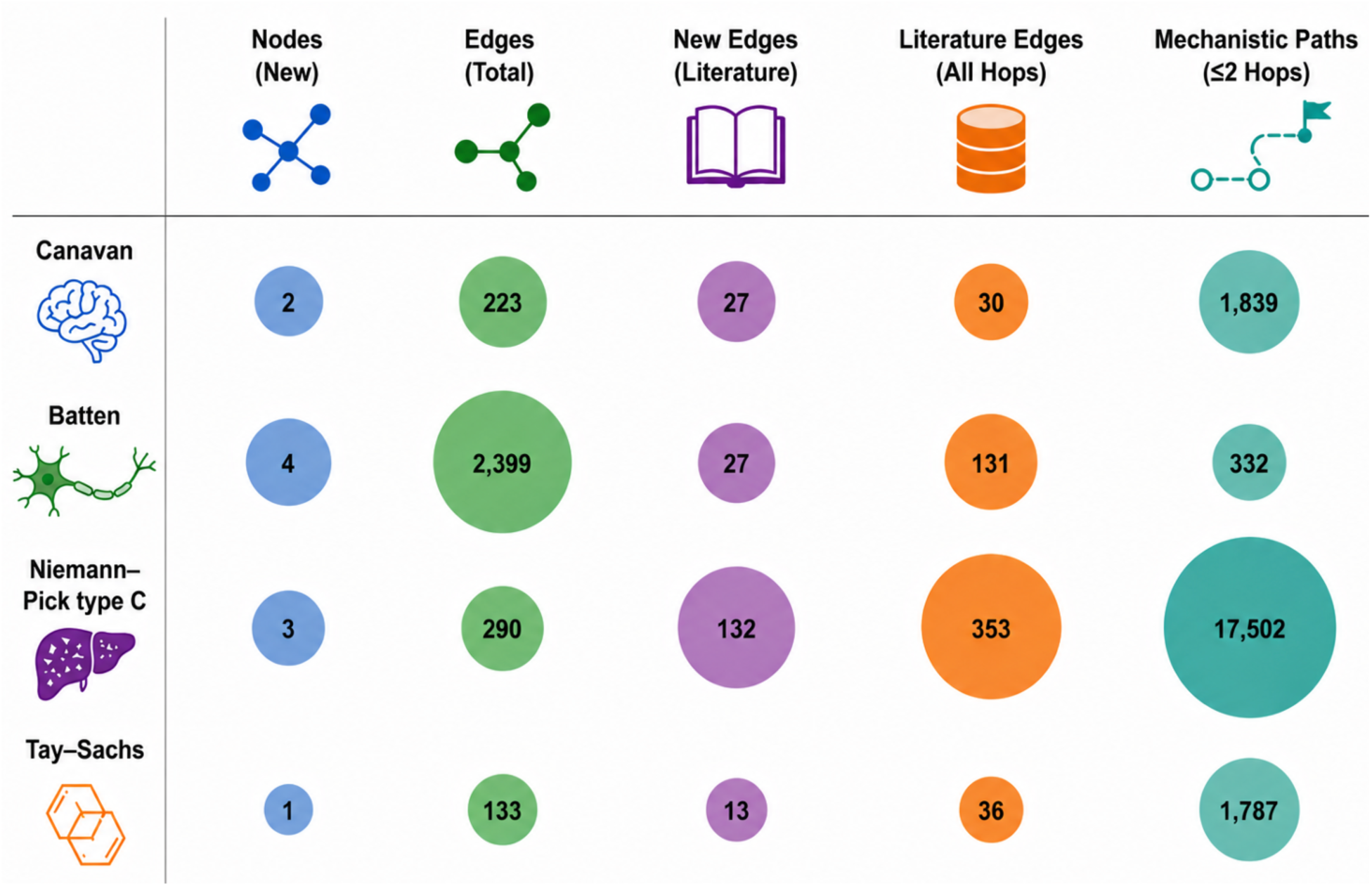
Graph enrichment around rare-disease anchors in PrimeKG-Plus vs Original PrimeKG.

Most new edges at one hop originate from the updated public databases (disease–protein and disease–phenotype links) and the literature integration contributes a smaller but targeted fraction. Mechanistic paths of length 1–6 hops passing through at least one literature edge were also enumerated per disease, with representative findings including core disease genes (ASPA, NPC1/NPC2, MFSD8, HEXA), candidate therapies (Miglustat, Arimoclomol for NPC), and cross-disease bridges (e.g., infantile NCL → Tay–Sachs via APOE). These path summaries support new hypothesis generation and expert-reviewed mechanistic claims that Original PrimeKG cannot recover.

## Conclusions

Taken together, existing biomedical knowledge graphs remain constrained by temporal staleness, limited rare-disease coverage, and challenges in reproducibility. PrimeKG-Plus addresses these limitations by combining synchronized biomedical database updates with curated rare-disease literature in a fully reproducible, PrimeKG-compatible resource. By restoring the temporal relevance of PrimeKG while expanding rare-disease knowledge, it provides an up-to-date foundation for drug repurposing, rare-disease research, and downstream machine-learning applications.

## Availability of source code and requirements

### PrimeKG-Plus

Project name: PrimeKG-Plus

- Project homepage: https://github.com/DSDD-UCPH/PrimeKG-Plus
- License: Software code under MIT license; curated literature-derived relations under CC-BY 4.0; integrated third-party biomedical resources remain subject to their respective licenses
- Operating system(s): Platform independent
- Programming language: Python
- Other requirements: pandas and Jupyter for graph assembly; licensed upstream dumps (e.g., DrugBank, UMLS) must be obtained from the original providers for full rebuilds
- PrimeKG-Plus code and knowledge-graph builds are also available through Zenodo (DOI: 10.5281/zenodo.20796545).

### Data availability

All knowledge-graph edge lists and node tables follow the 12-column PrimeKG schema and are provided in CSV format, together with literature-derived relation tables, curated entity-mapping files, and auxiliary rebuild outputs. PrimeKG-Plus is released as two graph builds: `primekg_plus.csv`, the base knowledge graph built from updated public biomedical databases only (no PubMed-derived edges), and `primekg_plus_rd.csv`, the literature-augmented build for the four studied disorders (Canavan, Niemann–Pick type C, Tay–Sachs, and Batten), which contains every edge in `primekg_plus.csv` plus expert-validated relations extracted from the curated literature corpus.

Following the PrimeKG release model, all files are deposited in Zenodo (DOI: 10.5281/zenodo.20796545), with rebuild code, variable names, metadata, and per-file documentation available on GitHub (https://github.com/DSDD-UCPH/PrimeKG-Plus); the release layout is shown in Figure 4. Raw upstream database dumps (e.g., DrugBank, UMLS) are not redistributed for licensing and size reasons and can be retrieved from the original providers using the documented primary_data_prep/ and additional_data_source/ workflows.

## Additional files

Additional file 1. Supplementary Information. Supplementary tables (S1–S11) — including literature-curation templates, schema definitions (Supplementary Information Section A; Tables S5–S8), audit files, and additional validation outputs — together with Supplementary Figures S1– S3. These materials accompany the manuscript and are also included in the Zenodo and GitHub release bundle.

## List of abbreviations

AI: artificial intelligence
API: application programming interface
ATC: Anatomical Therapeutic Chemical Classification
BERT: Bidirectional Encoder Representations from Transformers
BP: biological process
CC: cellular component
ChEMBL: Chemical database of EMBL
CSV: comma-separated values
CTD: Comparative Toxicogenomics Database
CUI: Concept Unique Identifier
DisGeNET: Disease–Gene Network
DOI: digital object identifier
FAIR: Findable, Accessible, Interoperable, and Reusable
FDA: U.S. Food and Drug Administration
GO: Gene Ontology
HGNC: HUGO Gene Nomenclature Committee
HPO: Human Phenotype Ontology
KG: knowledge graph
MedDRA: Medical Dictionary for Regulatory Activities
MeSH: Medical Subject Headings
MF: molecular function
MONDO: Mondo Disease Ontology
MSH: Medical Subject Headings (UMLS source abbreviation)
NCBI: National Center for Biotechnology Information
NCI: National Cancer Institute Thesaurus
NME: new molecular entity
OMIM: Online Mendelian Inheritance in Man
ORDO: Orphanet Rare Disease Ontology
PMC: PubMed Central
PrimeKG: Precision Medicine Knowledge Graph
PT: preferred term
QC: quality control
REST: Representational State Transfer
RRID: Research Resource Identifier
SapBERT: Self-alignment Pretrained BERT
SBERT: Sentence-BERT
SIDER: Side Effect Resource
SNOMEDCT_US: SNOMED CT US Edition
SQL: Structured Query Language
STRING: Search Tool for the Retrieval of Interacting Genes/Proteins
TIAB: Title/Abstract
TUI: Type Unique Identifier
UBERON: Uber-anatomy ontology
UMLS: Unified Medical Language System
XML: Extensible Markup Language.

## Funding

Lundbeck Foundation [R449-2023-1394]. Funding for open access charge: Lundbeck Foundation.

## Competing interests

A.J.K. is an employee and shareholder of Kvantify. A.J.K. is co-founder and CTO of Synamics Therapeutics.

## Author Contributions

Conceptualization (TTDN, AJK, ZT, DTN), Methodology (All authors), Software (TTDN), Validation (TTDN, TNP, ASH, ZT, DTN, AJK), Investigation (TTDN, ZT, DTN, TNP), Data Curation (TTDN, TNP, QHN, HDLP, BANL, AMA, PNT, NNC), Writing – Original Draft (TTDN), Writing – Review and Editing (All authors), Visualization (TTDN), Supervision (AJK), Funding Acquisition (TTDN, AJK, DTN, ASH)

## Acknowledgements

The authors would like to thank all student participants for their enthusiastic participation. We’d also like to thank Hue Phuong Tran, Ha Khue Nguyen, Vy Nguyen Dinh, Minh-Hieu Hoang Ngoc who have assisted in data curation.

## Notes

https://doi.org/10.5281/zenodo.20796545

https://github.com/DSDD-UCPH/PrimeKG-Plus

### Summary of Updates

The title was adjusted to emphasize a rare disease focus The content was re-structured but no new information was added

