## Supplementary information for "PrimeKG-Plus: a refreshed and rare-disease-enriched precision medicine knowledge graph"

### PrimeKG-Plus: a literature-derived expansion of a multimodal precision medicine knowledge graph

#### Outline:

Part I — Comparison of Original PrimeKG and PrimeKG-Plus (Tables S1–S4).

Part II — Supplementary Methods (Sections A, B, C, D, and E; Tables S5–S11; Figures S1–S3).

#### Part I. Comparison of Original PrimeKG and PrimeKG-Plus

Supplementary Tables S1–S4 summarize structural differences between the published Original PrimeKG (no_dup_kg.csv; Chandak et al.) and PrimeKG-Plus (20260529-primekg_plus.csv). All edge counts are directed unless otherwise noted. Tables S1–S2 include a plain-language reason for each change (intentional filtering, source updates, or ontology refresh).

##### Table S1. Node counts by entity type

*Total nodes: Original PrimeKG = 129,375; PrimeKG-Plus = 129,317.*

| Entity type | Original (n) | Original (%) | PrimeKG-Plus (n) | PrimeKG-Plus (%) | Reason for change |
|---|---|---|---|---|---|
| biological_process | 28,642 | 22.1 | 25,153 | 19.5 | Updated GO release; obsolete biological-process terms removed during harmonization. |
| gene/protein | 27,671 | 21.4 | 23,668 | 18.3 | Stricter protein ID |

| | | | | | |
|---|---|---|---|---|---|
| | | | | | harmonization and removal of unresolved Entrez mappings in updated sources. |
| disease | 17,080 | 13.2 | 19,884 | 15.4 | MONDO 2025 disease-only filter plus refined disease grouping (more singleton MONDO nodes, fewer grouped hubs). |
| effect/phenotype | 15,311 | 11.8 | 18,688 | 14.5 | Expanded HPO / MONDO–phenotype cross-references in the updated build. |
| anatomy | 14,035 | 10.8 | 14,611 | 11.3 | Additional UBERON anatomy terms from updated Bgee processing. |

| | | | | | |
|---|---|---|---|---|---|
| molecular_function | 11,169 | 8.6 | 10,143 | 7.8 | Updated GO molecular-function release; obsolete terms excluded. |
| drug | 7,957 | 6.2 | 9,384 | 7.3 | DrugBank 5.1.13 vocabulary update and broader drug node coverage. |
| cellular_component | 4,176 | 3.2 | 4,058 | 3.1 | Minor GO cellular-component pruning; net change small. |
| pathway | 2,516 | 1.9 | 2,848 | 2.2 | Reactome 2024 update added pathway nodes. |
| exposure | 818 | 0.6 | 880 | 0.7 | Updated CTD exposure processing (October 2025). |

| | **Original (n)** | **PrimeKG-Plus (n)** | **Reason for change** |
|---|---|---|---|
| Total | 129,375 | 129,317 | Net −2.0% nodes: disease/phenotype/drug up; gene/GO and |

| | | | |
|---|---|---|---|
| | | | anatomy-expression filtering down (see row notes). |

**Table S2. Directed edge counts by relation type**

*Total directed edges: Original PrimeKG = 8,097,472; PrimeKG-Plus = 7,683,206.*

| Relation type | Original (n) | Original (%) | PrimeKG-Plus (n) | PrimeKG-Plus (%) | Reason for change |
|---|---|---|---|---|---|
| anatomy_protein_present | 3,033,804 | 37.5 | 1,860,930 | 24.2 | Bgee reprocessed with expression_rank ≤ 5,000 (vs all ranks in Original PrimeKG), reducing redundant anatomy–gene edges. |
| drug_drug | 2,672,628 | 33.0 | 2,855,310 | 37.2 | Updated drug–drug interaction sources in DrugBank / supplementary drug interaction tables. |
| protein_protein | 642,150 | 7.9 | 765,720 | 10.0 | STRING 12.0 high-confidence human PPIs |

| | | | | | |
|---|---|---|---|---|---|
| | | | | | added to legacy PrimeKG pairs. |
| disease_phenotype_positive | 300,629 | 3.7 | 369,608 | 4.8 | Updated MONDO–HPO phenotype cross-references. |
| bioprocess_protein | 289,599 | 3.6 | 320,402 | 4.2 | Updated NCBI gene2go / GO annotation release. |
| cellcomp_protein | 166,782 | 2.1 | 192,902 | 2.5 | Updated gene–cellular-component annotations. |
| disease_protein | 160,813 | 2.0 | 516,740 | 6.7 | Updated DisGeNET plus newly integrated OpenTargets disease–gene associations; disease grouping aggregates member edges onto MONDO_grouped hubs. |
| molfunc_protein | 139,053 | 1.7 | 186,978 | 2.4 | Updated gene–molecular-function annotations |

| | | | | | |
|---|---|---|---|---|---|
| | | | | | from GO 2025-10-10. |
| drug_effect | 129,568 | 1.6 | 121,840 | 1.6 | SIDER + nSIDES integration refresh; modest net decrease after deduplication. |
| bioprocess_bioprocess | 105,772 | 1.3 | 85,940 | 1.1 | GO hierarchy update; obsolete biological-process is_a edges removed. |
| pathway_protein | 85,292 | 1.1 | 94,046 | 1.2 | Reactome 2024 pathway–protein mapping update. |
| disease_disease | 64,388 | 0.8 | 67,720 | 0.9 | MONDO hierarchy refresh; some parent–child edges removed as obsolete. |
| contraindication | 61,350 | 0.8 | 26,826 | 0.3 | Same MONDO / UMLS mapping policy as indications; ambiguous drug–disease links dropped. |

| | | | | | |
|---|---|---|---|---|---|
| drug_protein | 50,936 | 0.6 | 62,948 | 0.8 | DrugBank 5.1.13 — more carrier, enzyme, target, and transporter links. |
| anatomy_protein_absent | 39,774 | 0.5 | 1,476 | 0.0 | Same Bgee expression filter; absent calls largely dropped with stricter threshold. |
| phenotype_phenotype | 37,472 | 0.5 | 44,358 | 0.6 | HPO hierarchy update added phenotype–phenotype edges. |
| anatomy_anatomy | 28,064 | 0.3 | 29,212 | 0.4 | UBERON anatomy hierarchy update. |
| molfunc_molfunc | 27,148 | 0.3 | 24,950 | 0.3 | GO hierarchy update; obsolete molecular-function is_a edges removed. |
| indication | 18,776 | 0.2 | 10,006 | 0.1 | Stricter MONDO disease curation and |

| | | | | | |
|---|---|---|---|---|---|
| | | | | | UMLS→MONDO single-CUI mapping; updated DrugCentral integration. |
| cellcomp_cellcomp | 9,690 | 0.1 | 9,302 | 0.1 | GO hierarchy update; minor pruning of cellular-component is_a edges. |
| phenotype_protein | 6,660 | 0.1 | 6,620 | 0.1 | Stable HPO–protein mapping; near-unchanged after deduplication. |
| off-label use | 5,136 | 0.1 | 2,398 | 0.0 | Same MONDO / UMLS mapping policy as indications. |
| pathway_pathway | 5,070 | 0.1 | 5,728 | 0.1 | Reactome pathway hierarchy update. |
| exposure_disease | 4,608 | 0.1 | 4,770 | 0.1 | Updated CTD exposure–disease events with MONDO remap. |

| | | | | | |
|---|---|---|---|---|---|
| exposure_exposure | 4,140 | 0.1 | 4,884 | 0.1 | CTD exposure hierarchy update. |
| exposure_bioprocess | 3,250 | 0.0 | 4,206 | 0.1 | Additional CTD exposure–biological-process links. |
| exposure_protein | 2,424 | 0.0 | 6,086 | 0.1 | Expanded CTD exposure–protein associations in 2025 processing. |
| disease_phenotype_negative | 2,386 | 0.0 | 1,180 | 0.0 | Fewer negative phenotype links after MONDO/HPO remap and filtering. |
| exposure_molfunc | 90 | 0.0 | 94 | 0.0 | Minor CTD exposure–molecular-function update. |
| exposure_cellcomp | 20 | 0.0 | 26 | 0.0 | Minor CTD exposure–cellular-component update. |

| | Original (n) | PrimeKG-Plus (n) | Reason for change |
|---|---|---|---|

| | | | |
|---|---|---|---|
| Total | 8,097,472 | 7,683,206 | Net −3.3% edges: fewer anatomy/drug–disease edges (stricter filters); more disease–gene and PPI edges. |

**Table S3. Disease node grouping statistics**

| **Graph** | **Total disease nodes** | **MONDO** | **MONDO_grouped** | **Mean group size** | **Median** | **Max** | **Groups >10 members** |
|---|---|---|---|---|---|---|---|
| Original PrimeKG | 17,080 | 15,813 | 1,267 | 5 | 3 | 87 | 106 |
| PrimeKG-Plus | 19,884 | 18,704 | 1,180 | 6.5 | 4 | 116 | 141 |

**Table S4. Disease-node degree distribution**

| **Disease subset** | **n** | **Mean degree** | **Median** | **Min** | **Max** |
|---|---|---|---|---|---|
| Original — all diseases | 17,080 | 40 | 12 | 2 | 3,048 |
| Original — MONDO | 15,813 | 33.9 | 10 | 2 | 3,048 |
| Original — MONDO_grouped | 1,267 | 115.1 | 66 | 2 | 1,952 |
| PrimeKG-Plus — all diseases | 19,884 | 53.7 | 8 | 2 | 10,534 |

| | | | | | |
|---|---|---|---|---|---|
| PrimeKG-Plus — MONDO | 18,704 | 43.5 | 8 | 2 | 10,534 |
| PrimeKG-Plus — MONDO_grouped | 1,180 | 214.4 | 116 | 2 | 4,552 |

**Notes**

- Original PrimeKG: published graph (no_dup_kg.csv) from the PrimeKG dataverse release.
- PrimeKG-Plus: build dated 20260529, incorporating updated source releases, UMLS–MONDO no-multi-mapping curation, OpenTargets disease–protein associations, and BGEE anatomy expression filtering (expression_rank ≤ 5000)
- Disease grouping merges synonym/similar MONDO disease nodes into MONDO_grouped hub nodes, reducing redundant edges while increasing hub degree (Table S4).

### Part II — Supplementary Methods

#### Section A. Literature curation prompt

##### Prompt

Read and understand the content in the file [ABSTRACT + INTRODUCTION] and construct a data table according to the guidelines provided in the file Curation Requirements.md.docx.

**Table S5. Literature curation output columns.**

| # | Column name | Description |
|---|---|---|
| 1 | doi/PMID | — |
| 2 | Abstract/Introduction | — |
| 3 | journal type | See options below |
| 4 | experiment | See options below |
| 5 | model | See options below |
| 6 | relation | Per Curation_Requirements.md |

| | | |
|---|---|---|
| 7 | display_relation | Per Curation_Requirements.md |
| 8 | x_name | Per Curation_Requirements.md |
| 9 | x_type | Per Curation_Requirements.md |
| 10 | y_name | Per Curation_Requirements.md |
| 11 | y_type | Per Curation_Requirements.md |
| 12 | note | — |

**Column 3 (journal type): select one of — "case report", "original article", "review", or "case report + review".**

Column 4 (experiment): select one of — "in vivo", "in vitro", or "ex vivo", depending on the experimental context.

Column 5 (model): select one of — "human", "mice", "cell line", or "human and mice", depending on the biological model.

Columns 6, 7, 8, and 9: fill according to Curation_Requirements.md.

#### 2. Content of Curation_Requirements.md

For each abstract, extract biomedical relationships between entities. Each relationship must belong to one of the predefined types below.

##### 2.1 Relationship types

**Table S6. Predefined PrimeKG relationship types.**

| Relation type | Description |
|---|---|
| anatomy_anatomy | is_a between anatomical structures |
| anatomy_protein_absent | protein absent in an anatomical structure |
| anatomy_protein_present | protein present or expressed in an anatomical structure |
| bioprocess_bioprocess | is_a between biological processes |
| bioprocess_protein | protein participates in a biological process |
| cellcomp_cellcomp | is_a between cellular components |
| cellcomp_protein | protein located in or associated with a cellular component |

| | |
|---|---|
| contraindication | disease where a drug is advised against |
| disease_disease | is_a relationship between diseases |
| disease_phenotype_negative | disease linked to absence of a phenotype |
| disease_phenotype_positive | disease linked to presence of a phenotype |
| disease_protein | disease linked to a protein (mutation, biomarker, dysregulation) |
| drug_drug | pharmacokinetic/pharmacodynamic interaction, synergy, antagonism |
| drug_effect | drug causes an observable effect or phenotype |
| drug_protein | drug interacts with a protein (target/enzyme/transporter/carrier) |
| exposure_bioprocess | exposure affects biological process |
| exposure_cellcomp | exposure affects cellular component |
| exposure_disease | exposure linked to disease |
| exposure_exposure | is_a between exposures |
| exposure_molfunc | exposure affects molecular function |
| exposure_protein | exposure affects protein activity or levels |
| indication | disease that a drug treats or prevents |
| molfunc_molfunc | is_a between molecular functions |
| molfunc_protein | protein has molecular function |
| off_label_use | non-approved indication |
| pathway_pathway | is_a between pathways |
| pathway_protein | protein acts in a pathway |
| phenotype_phenotype | is_a or association between phenotypes |
| phenotype_protein | phenotype associated with or influenced by a protein |
| protein_protein | protein–protein interaction/regulation |

#### 2.2 Entity types

The two entities in each relationship must be one of: disease, protein/gene, drug, anatomy, phenotype, biological_process, cellular_component, molecular_function, pathway, exposure, pathology.

##### 2.3 Example rows

**Table S7. Example curated relationship rows.**

| relation | Display relation | x_name | x_type | y_name | y_type | note |
|---|---|---|---|---|---|---|
| protein_protein | ppi | PHYHIP | gene/protein | KIF15 | gene/protein | x_name: synonym |
| protein_protein | ppi | GPANK1 | gene/protein | PNMA1 | gene/protein | your note |
| drug_protein | carrier | Flurandrenolide | drug | SERPINA6 | gene/protein | your notes |
| drug_protein | carrier | Prednisolone | drug | SERPINA6 | gene/protein | … |
| phenotype_protein | assoc_with | PRSS35 | gene/protein | Hepatomegaly | phenotype | … |

##### 2.4 Column definitions

**Table S8. Column definitions for curated relationships.**

| Column | Meaning |
|---|---|
| relation | The relationship type. |
| display_relation | Usually identical to relation; in some cases must be specified in more detail (see Notes). |
| x_name | Name of the first entity. |
| x_type | Entity type of the first entity. |
| y_name | Name of the second entity. |
| y_type | Entity type of the second entity. |

**Notes**

1. If relation = "drug_protein", display_relation must be one of: 'carrier', 'enzyme', 'target', or 'transporter'.
2. If you identify any potentially new or additional relationship types that may be useful, record them in the note column.

**Figure S1. Curation team roles and quality-control workflow.**

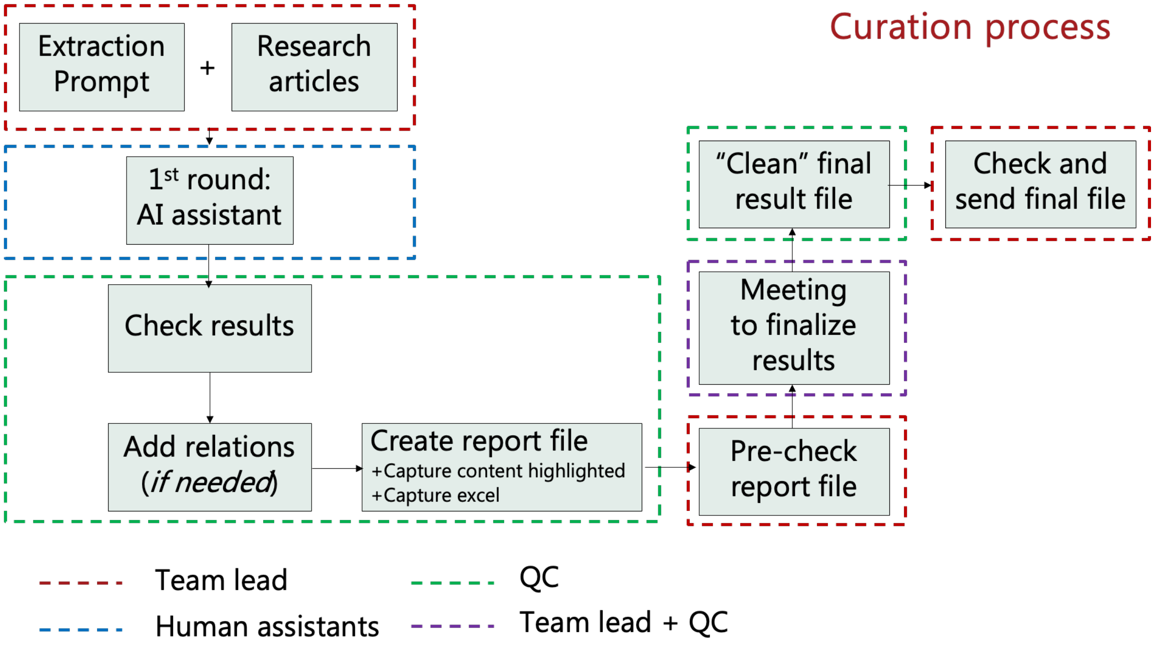

**Figure S2. Article-level quality-control and correction workflow.**

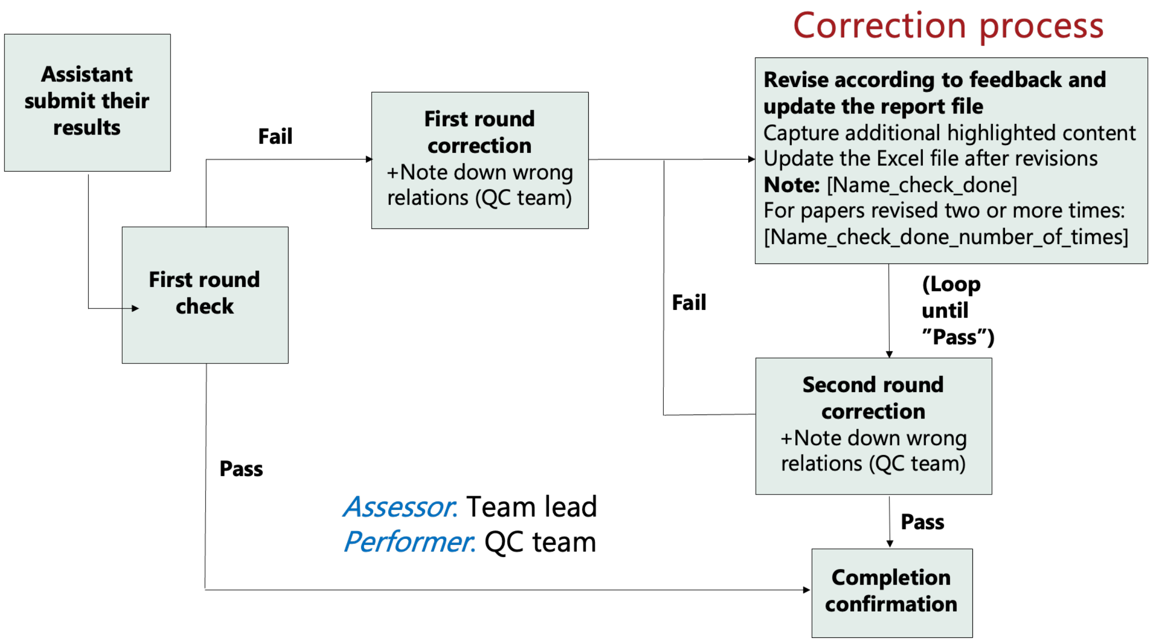


### Section B. UMLS semantic type validation

**Table S9. UMLS Semantic Types (TUIs) Accepted per Entity Type**

| Entity Type | TUI | Semantic Type Name | Notes |
|---|---|---|---|
| disease | T047 | Disease or Syndrome | Primary disease category |
| | T191 | Neoplastic Process | Tumor and cancer-related |
| | T048 | Mental or Behavioral Dysfunction | Psychiatric disease-related |
| gene/protein | T028 | Gene or Genome | Primary gene category |
| | T192 | Receptor | Treated as protein |
| drug | T121 | Pharmacologic Substance | Primary drug category |
| | T200 | Clinical Drug | Formulated drug products |
| | T195 | Antibiotic | Subset of pharmacologic substance |
| phenotype | T033 | Finding | Clinical findings (e.g., hypotonia) |
| | T034 | Laboratory or Test Result | Lab-based phenotypes |

| | | | |
|---|---|---|---|
| | T041 | Mental Process | Cognitive/behavioral phenotypes |
| | T184 | Sign or Symptom | Observable clinical signs |
| | T019 | Congenital Abnormality | Structural birth defects |
| | T020 | Acquired Abnormality | Post-natal structural changes |
| biological_process | T038 | Biologic Function | Broad biological processes |
| | T039 | Physiologic Function | Normal physiological processes |
| molecular_function | T043 | Cell Function | Cellular-level molecular functions |
| cellular_component | — | — | No direct TUI mapping available |
| pathway | T038 | Biologic Function | Closest approximation; no pathway-specific TUI |
| anatomy | T017 | Anatomical Structure | General anatomical entities |
| | T018 | Embryonic Structure | Developmental anatomy |
| | T021 | Fully Formed Anatomical Structure | Mature anatomical entities |
| | T022 | Body System | Organ systems |
| | T023 | Body Part, Organ, or Organ Component | Organs and substructures |
| | T024 | Tissue | Tissue-level structures |
| | T029 | Body Location or Region | Spatial body regions |
| | T030 | Body Space or Junction | Cavities and junctions |
| exposure | T060 | Diagnostic Procedure | Imaging and diagnostic tests (e.g., MRI) |
| | T061 | Therapeutic or Preventive Procedure | Clinical interventions |
| | T063 | Molecular Biology Research Technique | Laboratory research methods |
| pathology | T046 | Pathologic Function | Pathological processes (e.g., inflammation) |

#### Section C. OpenTargets score threshold justification

DisGeNET and Open Targets employ fundamentally different scoring frameworks — DisGeNET assigns fixed per-source weights yielding a normalized confidence score (0–1), whereas Open Targets computes a hierarchical harmonic-sum aggregation across evidence types — making direct threshold equivalence inappropriate. To identify a threshold that retains associations of comparable evidence strength to DisGeNET (≥0.3), we compared score distributions under two OpenTargets cutoffs. At ≥0.1, the score distributions of overlapping DisGeNET–OpenTargets associations were well-aligned, indicating comparable evidence quality across both sources (Supplementary Figure S3, top row). At ≥0.3, OpenTargets distributions diverged markedly, excluding a large fraction of well-supported associations (Supplementary Figure S3, bottom row). We therefore retained direct target–disease associations from Open Targets (version 25.12.0) with an overall association score >0.1

**Figure S3. Score distribution for protein-disease associations in DisGeNet and OpenTarget**

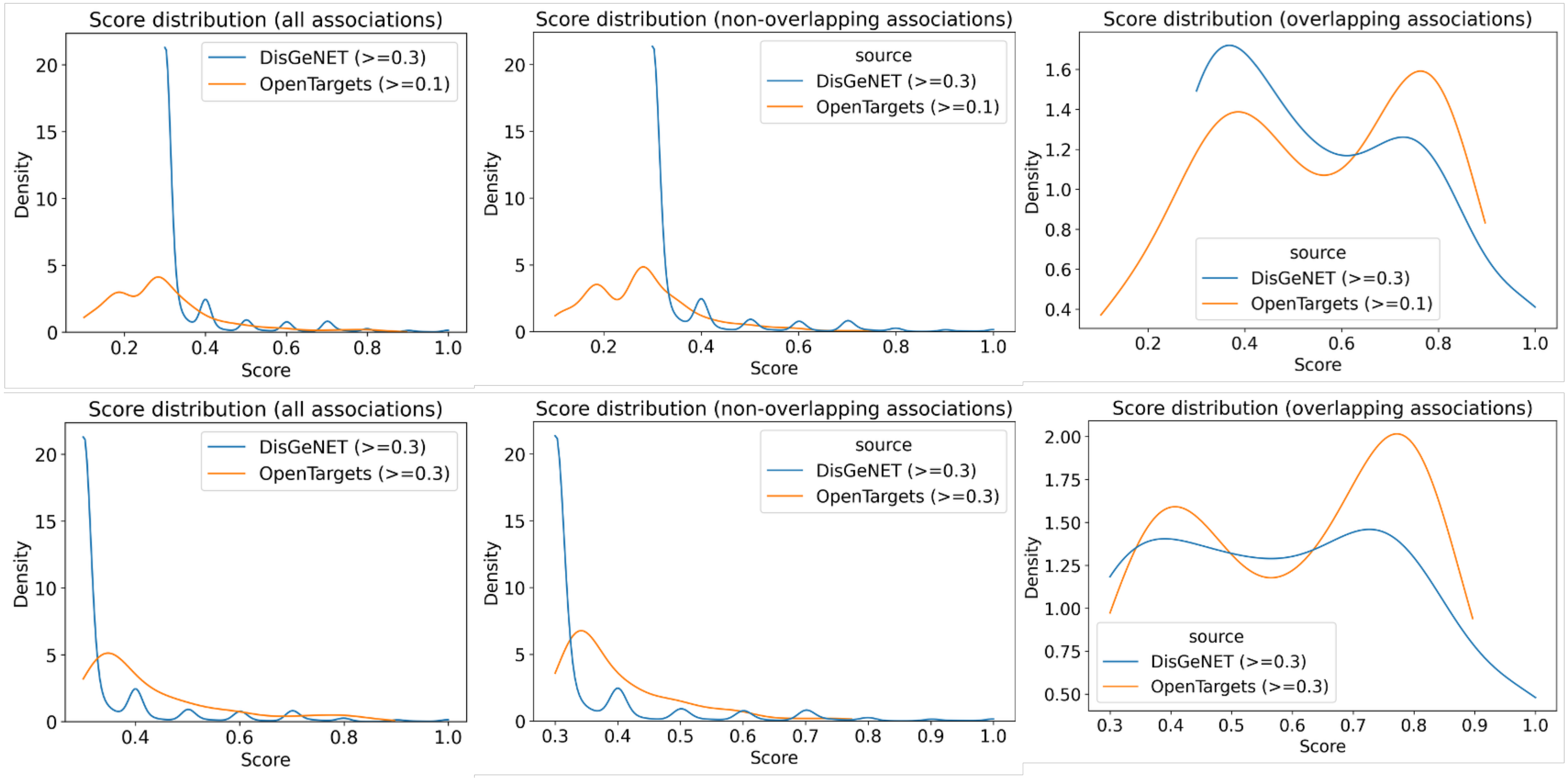

#### Section D. UMLS–MONDO bijective vocabulary

PrimeKG-Plus maps external disease identifiers (DrugCentral, DisGeNET, CTD, and others) to MONDO nodes via a curated UMLS–MONDO cross-reference table. The vocabulary used in published PrimeKG is not bijective: a single UMLS CUI can map to multiple MONDO IDs and vice versa, causing ambiguous rows to propagate as duplicated disease-centered edges during graph assembly (inner joins).

The original vocabulary contained 31,683 CUI–MONDO pairs; 10.35% of unique CUIs mapped to more than one MONDO ID, and 20.17% of unique MONDO IDs mapped to more than one CUI. Overall, 49.28% of pair rows participated in at least one one-to-many association (Supplementary Table S11).

To prevent spurious edge duplication, PrimeKG-Plus uses a bijective vocabulary (21,347 pairs; v1) retaining only MONDO concepts with a single unambiguous UMLS cross-reference in the Monarch Initiative database. An extended curation pass adding UMLS Terminology Services API resolution recovers 10 additional pairs (v2; 21,357 pairs). Both versions eliminate multi-mapping entirely. A replay of the DrugCentral pipeline confirms the practical impact: the original vocabulary yields 31,118 rows versus 20,816 with the bijective file (−33.1%), attributable to mapping multiplicity rather than loss of source records.

Representative examples are provided in Supplementary Table S10.

**Table S10. Examples of mapping ambiguity resolved in PrimeKG-Plus.**

| Pattern | Identifier | Original mapping | Curated (bijective) |
|---|---|---|---|
| One CUI → many MONDO | UMLS C0019829 | 5 MONDO diseases (e.g., MONDO:9348 classic Hodgkin lymphoma; MONDO:4952 | MONDO:9348 only → 28 vs 140 pipeline rows; e.g. mechlorethamine: |

| | | | |
|---|---|---|---|
| | | Hodgkins lymphoma; MONDO:4604 Hodgkin's lymphoma, lymphocytic-histiocytic predominance) | 1 DC row → 5 indication edges |
| One CUI → two MONDO | UMLS C0001125 | MONDO:6040 (lactic acidosis) + MONDO:24306 (acquired lactic acidosis) | MONDO:6040 only → 23 vs 46 pipeline rows; e.g. metformin: 1 contraindication → 2 disease edges |
| One MONDO → many UMLS | MONDO:18076 tuberculosis | 49 UMLS CUIs (e.g., C0041296, C0026926, C0151332) | 1 preferred CUI per MONDO |
| One MONDO → two UMLS | MONDO:9348 classic Hodgkin lymphoma | C0019829 and C1333064 | C0019829 retained |

**Table S11. Summary of UMLS–MONDO vocabulary versions used in PrimeKG-Plus.**

| **Metric** | **Original vocabulary (umls_mondo.csv)** | **Curated bijective vocabulary (no_multi_mapping)** |
|---|---|---|
| CUI–MONDO pair rows | 31,683 | 21,347 (v1) / 21,357 (v2) |
| Unique UMLS CUIs | 28,349 | 21,347 / 21,357 |
| Unique MONDO IDs | 22,418 | 21,347 / 21,357 |

| | | |
|---|---|---|
| CUIs mapping to >1 MONDO | 2,935 (10.35% of CUIs) | 0 (0%) |
| MONDO IDs mapping to >1 CUI | 4,521 (20.17% of MONDO) | 0 (0%) |
| Pair rows in any one-to-many association | 15,613 (49.28%) | 0 (0%) |

#### Section E. Deviations from original PrimeKG scripts

##### 1. Refined disease grouping via improved subtype matching

The original grouping algorithm identified disease subtypes by stripping the final token of each disease name and comparing the resulting prefix to other entries. While effective for simple cases (e.g., "Gaucher disease type 1" → prefix "Gaucher disease type"), this approach produced two systematic failure modes: (i) names containing trailing punctuation (e.g., "Gaucher disease, type 1") were not matched due to the residual comma after suffix removal, and (ii) disease names without a subtype suffix (e.g., "Gaucher (disease)") were never passed to the fuzzy word-overlap check, causing them to be silently excluded from their disease group.

The revised algorithm addresses both issues by introducing a structured three-case matching function. First, exact string equality is tested as a fast-path check. Second, prefix containment via startswith handles punctuation-adjacent suffixes. Third, names with a recognized subtype suffix are matched after suffix removal, while names without such a suffix are passed directly to the word-overlap function. Together, these changes reduce false negatives in disease grouping without altering the underlying grouping criteria.

**Script:** *scripts/02_disease_grouping.ipynb*

##### 2. Retrieval of protein–protein interactions from the published PrimeKG graph

The original PrimeKG build script for protein–protein interactions (PPIs) was not available in the public repository. PrimeKG-Plus therefore retrieves PPI edges directly from the published PrimeKG graph file, retaining all edges of relation type protein_protein. To expand PPI coverage, additional interactions were integrated from STRING v12.0, filtered to experimentally validated associations (combined score ≥ 900). Edges already present in the retrieved PrimeKG PPI set were deduplicated prior to integration.

**Script:** *scripts/01_build_graph.ipynb (loads primary_data_prep/data/ppi/protein_protein.csv, which combines retrieved PrimeKG PPI edges with STRING v12.0 additions)*

##### 3. Open Targets disease–protein associations

Original PrimeKG relied on DisGeNET for disease–gene associations. PrimeKG-Plus adds direct target–disease associations from Open Targets Platform release 25.12.0 (association_overall_direct), harmonized to gene symbols and UMLS CUIs, filtered against existing DisGeNET pairs, and merged with DisGeNET-derived disease_protein edges during graph assembly. Score-threshold justification is provided in Section C.

**Script:** *additional_data_source/opentarget/process_opentarget.ipynb → primary_data_prep/data/disgenet/OpenTarget/OpenTarget_disease_protein_associations.csv; consumed by scripts/01_build_graph.ipynb*

##### 4. RepurposeDrugs Phase-4 indication edges

Approved Phase-4 drug–disease candidate pairs from RepurposeDrugs were resolved to DrugCentral-compatible (CAS registry number, UMLS CUI) rows. Pairs already present in the DrugCentral indication table were removed by anti-join before merging novel indications into the drug–disease edge set.

**Script:** *additional_data_source/repurposed_drug/process_repurposed_drug.ipynb → primary_data_prep/data/repurposed_drug/RepurposedDrug_Indication.csv; merged in scripts/01_build_graph.ipynb*

##### 5. SIDER + Open nSIDES drug–adverse-effect integration

PrimeKG-Plus reproduces the SIDER 4.1 MedDRA Preferred Term pipeline and augments it with high-confidence ingredient–effect pairs from Open nSIDES (v3.1.0). nSIDES MedDRA effects are mapped to UMLS CUIs via the SIDER effect vocabulary; RxNorm ingredients are mapped to ATC codes through DrugBank XML, yielding a SIDER-compatible drug_effect table that replaces the SIDER-only input used in the original build.

**Script:** *additional_data_source/sider_nsides/build_sider.py and additional_data_source/sider_nsides/process_sider_nsides.ipynb → primary_data_prep/data/sider/sider_with_nsides.csv; consumed by scripts/01_build_graph.ipynb*

#### 6. UMLS–MONDO bijective vocabulary (multi-mapping resolution)

Published PrimeKG uses a non-bijective UMLS–MONDO cross-reference table in which one CUI can map to multiple MONDO IDs and vice versa, inflating drug–disease and protein–disease edge counts during inner joins (Section D; Supplementary Tables S10–S11). PrimeKG-Plus applies a curated bijective vocabulary (20260510-umls_mondo_no_multi_mapping_v2.csv; 21,357 MONDO–UMLS pairs) everywhere a source encodes disease as a UMLS CUI and the graph standardizes on MONDO, including DrugCentral, DisGeNET/Open Targets merges, CTD exposure events, and literature entity resolution.

**Script:** *scripts/01_build_graph.ipynb (loads primary_data_prep/data/vocab/umls_mondo_bijective.csv); scripts/literature_curation/lib/entity_resolver.py (CUI→MONDO lookup during literature integration)*

#### 7. Literature-derived relations: PrimeKG matching, SapBERT retrieval, and graph integration

Curated rare-disease relations undergo a staged mapping workflow before integration into primekg_plus_rd.csv. Entity strings are first matched against existing PrimeKG-Plus nodes by normalized name. Unmatched entities are mapped through UMLS (CUI assignment and semantic-type validation per Section B, Table S9). For second-pass name repair, candidate replacements in the PrimeKG entity pool are retrieved with SapBERT cosine similarity over a precomputed UMLS embedding memmap and reranked with Sentence-BERT (SBERT). Resolved endpoints are checked against the PrimeKG relation schema; unsupported relation types are logged and skipped. Valid rows are deduplicated against existing directed edges in primekg_plus.csv before append.

Script: scripts/literature_curation/lib/query_sapbert_rerank_sbert.py (SapBERT top-k + SBERT rerank); scripts/literature_curation/lib/sapbert_pool_encode.py and scripts/literature_curation/lib/sapbert_encode_primekg_style.py (SapBERT encoding utilities); scripts/literature_curation/lib/entity_resolver.py (PrimeKG node matching and CUI→ontology resolution); scripts/literature_curation/lib/relation_config.py (relation normalization and PrimeKG schema check); scripts/literature_curation/09_integrate_primekg_plus_rd.py (final integration and audit exports; alias integrate_primekg_plus_rd.py); scripts/literature_curation/03_map_curated_entities_canavan.ipynb through

06_map_curated_entities_tay_sachs.ipynb (per-disease entity mapping); 07_finalize_post_curation.ipynb (second-search QC); 08_merge_expert_post_curation.py (expert-merge additional relations)
